# Sensory History Shapes Contrastive Neural Geometry in LP/Pulvinar-Prefrontal Cortex Circuits

**DOI:** 10.1101/2024.11.16.623977

**Authors:** Yi Ning Leow, Arundhati Natesan, Alexandria Barlowe, Sofie Ährlund-Richter, Cindy Luo, Mehrdad Jazayeri, Mriganka Sur

## Abstract

Prior expectations guide attention and support perceptual filtering for efficient processing during decision-making. Here we show that during a visual discrimination task, mice use prior stimulus history to guide ongoing choices by estimating differences in evidence between consecutive trials (|ΔDir|). The thalamic lateral posterior (LP)/pulvinar nucleus provides robust inputs to the Anterior Cingulate Cortex (ACC), which has been implicated in selective attention and predictive processing, but the function of the LP-ACC projection is unknown. We found that optogenetic manipulations of LP-ACC axons disrupted animals’ ability to effectively estimate and use information across stimulus history, leading to |ΔDir|-dependent biases. Two-photon calcium imaging of LP-ACC axons revealed an engagement-dependent low-dimensional organization of stimuli along a curved manifold. This representation was scaled by |ΔDir| in a manner that emphasized greater deviations from prior evidence. Thus, our work identifies the LP-ACC pathway as essential for selecting and evaluating stimuli relative to prior evidence to guide decisions.

## Introduction

Our perceptual landscape is constantly shaped by prior expectations and ongoing goals, which exert powerful influence over attentional and perceptual processing (*1–4*). Such expectation-dependent modulation can bias how incoming sensory evidence is evaluated, prioritizing information that is relevant given recent experience or current goals. Beyond transient attentional gain, such influences may operate at the level of decision formation, shaping how sensory information is used to guide choice. The pulvinar, a higher order visual thalamic nucleus, is well positioned to support such computations through widespread reciprocal connectivity with visual, associative and frontal cortices. Mutual interactions between the pulvinar and the frontal-parietal attention network have been implicated in regulating sustained and selective attentional allocation during visual decision-making (*5–22*). Prior work established compelling evidence for the pulvinar’s contributions to visuospatial attention and filtering of ongoing distractors, particularly through interactions with visual cortex and parietal cortex (*11–22*). However, it remains unclear how pulvinar outputs to the frontal cortex contribute to decision-related processes beyond established roles in attention.

The lateral posterior nucleus (LP) is the rodent homolog of the primate pulvinar, and shares most of the key cortical partners (*23,24*), indicating that the pulvinar’s hub-like functions are conserved across species (*8,12,15*). Recent work suggests that LP outputs to different cortical areas could carry distinct visual and contextual information (*25*), underscoring a need for further target-specific functional dissection of information carried by LP outputs. Here, we focus on LP projections to the anterior cingulate cortex (ACC), a prefrontal region critical for action selection, error computations and anticipating cues that provide information and resolve uncertainty (*26–34*). Reciprocal LP-ACC circuits in rodents integrate inputs from comparable cortical and subcortical regions as the primate pulvinar-frontal circuits (*21*). Moreover, relative to LP inputs to other visual areas, LP-ACC pathways preferentially integrate inputs from prefrontal and association areas (*23,24*), positioning this pathway to incorporate internal, history or reward-related signals alongside incoming sensory information. This raises the key question of how LP-ACC interactions influence the evaluation of sensory evidence during decision-making.

Here, we developed a behavioral and circuit-level approach to examine how prior and current evidence are integrated in LP/pulvinar projections to ACC. Using a visual discrimination task in mice, we asked how trial-to-trial differences in sensory evidence shape choice behavior and examined how LP-ACC inputs contribute causally to the process. Using optogenetic perturbations during the task performance and two-photon imaging of LP axons in ACC, we investigated how history-dependent signals are represented in this pathway and how its disruption alters decision-making. Together, our results identify a thalamocortical circuit that links trial history to the evaluation of ongoing sensory evidence, providing a contrastive signal that governs history-dependent evidence evaluation.

## Results

### Mice make choices based on estimated differences across consecutive stimuli

We trained mice on a two-choice visual discrimination task to test how trial-to-trial differences in sensory evidence influence perceptual decisions. Head-fixed mice learned to discriminate between two global dot movement directions (0°, temporally or 180°, nasally) in random dot kinematograms (RDK) presented for 1.5s, and reported their choice by licking either the left or right lick spout during a 1.5-s response window (Fig. 1A, B). Stimulus uncertainty was varied by presenting RDKs at graded levels of coherence (100%, 64%, 32%, or 16% coherence) with equal probabilities. The task was self-paced, requiring mice to withhold licking and run on a treadmill for 1s to initiate trials. Correct responses were rewarded with water, while errors were punished with a timeout.

**Figure 1:**
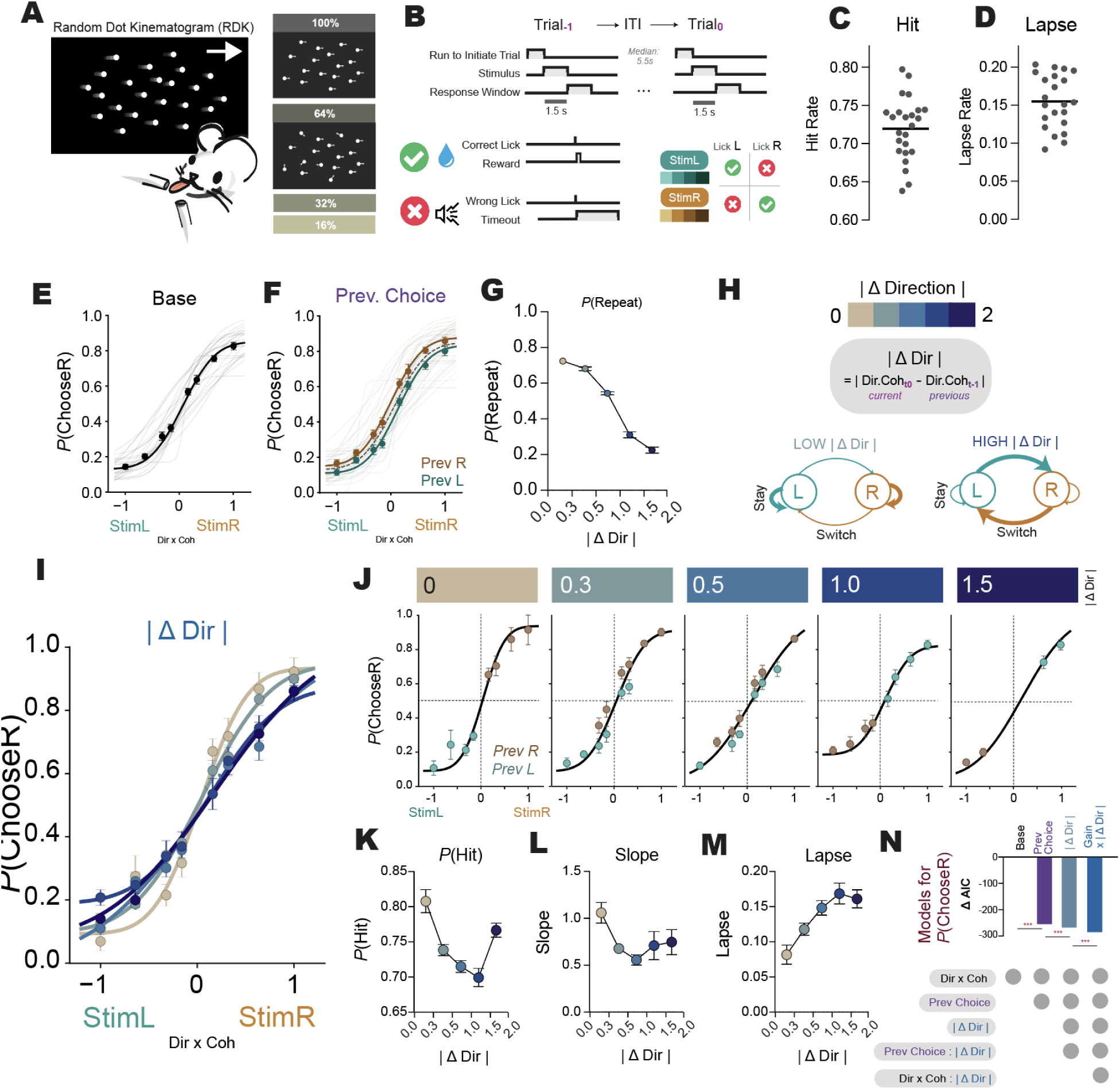
Mice make choices based on estimated differences across consecutive stimuli. (**A**) Schematic of the two-alternative choice random dot kinematogram (RDK) visual discrimination task. Head-fixed mice reported perceived motion direction by licking a left or right spout. Motion direction (0° Temporal / 180° Nasal) and coherence (16–100%) varied across trials. (**B**) Trial Structure and epochs. Mice run to initiate a trial and are presented with an RDK stimulus for 1.5 seconds. Mice report their choices by licking either the left or right spout during the 1.5 second response window. 100ms upon choice, correct choices are rewarded with water delivery while wrong choices are followed by a loud white noise and flashing screen. Successive trials are separated by a median inter-trial interval (ITI) of 5.5s. (**C**) Hit rates, sampling uniformly from all coherence levels. (**D**) Mean lapses. For (C) and (D) each dot represents a mouse, bold line indicates population mean across n = 24 mice. (**E**) Probability of choosing right (P(ChooseR)) as a function of signed stimulus strength (direction x coherence). Population mean in bold black. Gray lines indicate individual animals. n = 24 mice. (**F**) Psychometric functions conditioned on previous choice indicating choice history biases. (**G**) Probability of repeating the previous choice (P(Repeat)) as a function of the absolute change in stimulus direction between consecutive trials (|ΔDir|). (**H**) Schematic illustrating the definition of |ΔDir| as the absolute difference in signed stimulus direction between the current and previous trial, and its relationship to stay versus switch behavior. (**I**) Population-level psychometric functions stratified by |ΔDir|. N = 22 mice. (**J**) Psychometric functions separated by previous choice (PrevL vs PrevR) within each |ΔDir| bin. (**K** to **M**) (K) Hit rate (P(Hit)), (L) Psychometric slopes, and (M) Lapse rate, as a function of |ΔDir| (**N**) Best fit model (lowest AIC) for P(ChooseR) incorporated |ΔDir| interactions with previous choice terms and evidence. Full details of models in Table S3.

Trained mice showed coherence-dependent psychometric performance (Fig. 1C). While choice behavior was largely dependent on the current stimulus evidence, mice nevertheless showed serial choice biases, with a propensity to repeat a previously rewarded choice (Fig. 1D). Importantly, we found that the probability of repeating a choice was highly dependent on the absolute difference between signed coherences across trials, which we denote as |ΔDir| (Fig. 1E, F and figs. S1A-C). When the current and previous stimuli were more similar (low |ΔDir|), mice were more likely to repeat their previously rewarded choice. Conversely, when consecutive stimuli were most different (high |Δ Dir|), they were least likely to repeat. The opposite was observed following error trials, revealing an outcome-dependent use of |ΔDir| (figs. S1E and F).

This |ΔDir|-dependent strategy extends beyond classical win-stay-lose-switch strategies by explicitly comparing how similar current and previous evidence are. Notably, the probability of repeating a choice showed a steeper dependence on |ΔDir| than expected from simulated null models of mean psychometric performance that had no history dependence, or models with only simple serial choice biases independent of previous stimulus strength (figs. S1K-L). Thus, this relationship could not be fully explained by static choice biases, performance at different stimulus difficulties, or stimulus presentation order, but instead required explicit comparison of trial-by-trial differences.

Trial-to-trial sensory differences strongly modulated psychometric performance (Fig. 1G). Following correct trials, increasing |ΔDir| was associated with reduced hit rates (Fig. 1K), shallower psychometric slopes (Fig. 1L), and increased lapse rates (Fig. 1M), with little effect on decision threshold (fig. S1M). These effects were symmetric with respect to previous choice (Fig. 1H; figs. S2A–D). Notably, performance varied non-monotonically with |ΔDir| performance (Fig 1K and L), peaking when consecutive stimuli were similar and declining at intermediate values, consistent with a switch-related cost that is only offset when sensory differences become large. After errors, we observed the opposite trend from after rewarded trials – hit rates improved and lapses decreased with greater |ΔDir| (fig. S1G-J).

Trial-by-trial choice behavior was best fitted by models incorporating |ΔDir| interactions with evidence and previous choice, outperforming models that simply accounted for the previous choice (Fig. 1N, Table S3). Thus, mice performed best when consecutive stimuli were similar and when repeating their previously rewarded choices, and performed worse when they were required to switch. Increasing |ΔDir| also increased the probability of mice omitting the trial (fig. S1N) and was also associated with greater response latency (fig. S1O).

Together, these results implied that mice retained a memory of the previous stimulus estimate, and the comparison with the current stimulus facilitated decisions to switch or stay, based on the outcome of the previous choice. Such short-term memory could be expected to decay with time, and indeed |ΔDir| had a diminishing influence on the probability of repeating a choice with increasing inter-trial intervals (ITI) (fig. S1P to S).

This |ΔDir|-dependent strategy was evident across all stimulus coherences rather than selectively enhanced at low coherence; instead, |ΔDir|-dependent modulation of choice repetition was strongest for high-coherence stimuli, consistent with a comparison of reliable sensory estimates (fig. S2E), while its impact on performance was comparable across coherence levels (fig. S2F), reflecting variability in the accuracy consequences of switching across levels of sensory reliability. Taken together, our findings demonstrate that mice use relative evidence across consecutive trials to modulate classical win-stay-lose-switch strategies.

### Activating LP-ACC projections biases choice in a |ΔDir|-dependent manner

The comparison of sensory evidence across trials requires a circuit that integrates both current sensory input and recent evidence, to ultimately influence action selection. To test whether LP inputs to ACC causally contribute to this process, we optogenetically stimulated LP axons in ACC. LP-ACC axons expressing the light-activated opsin ChR2 unilaterally were activated by illuminating the ACC with blue light during visual stimulus presentation (Fig. 2A and B, fig. S3A). Visual stimuli were always presented on the right screen, enabling us to dissociate effects driven by contralateral visual input from effects on evidence-to-choice transformation.

**Figure 2:**
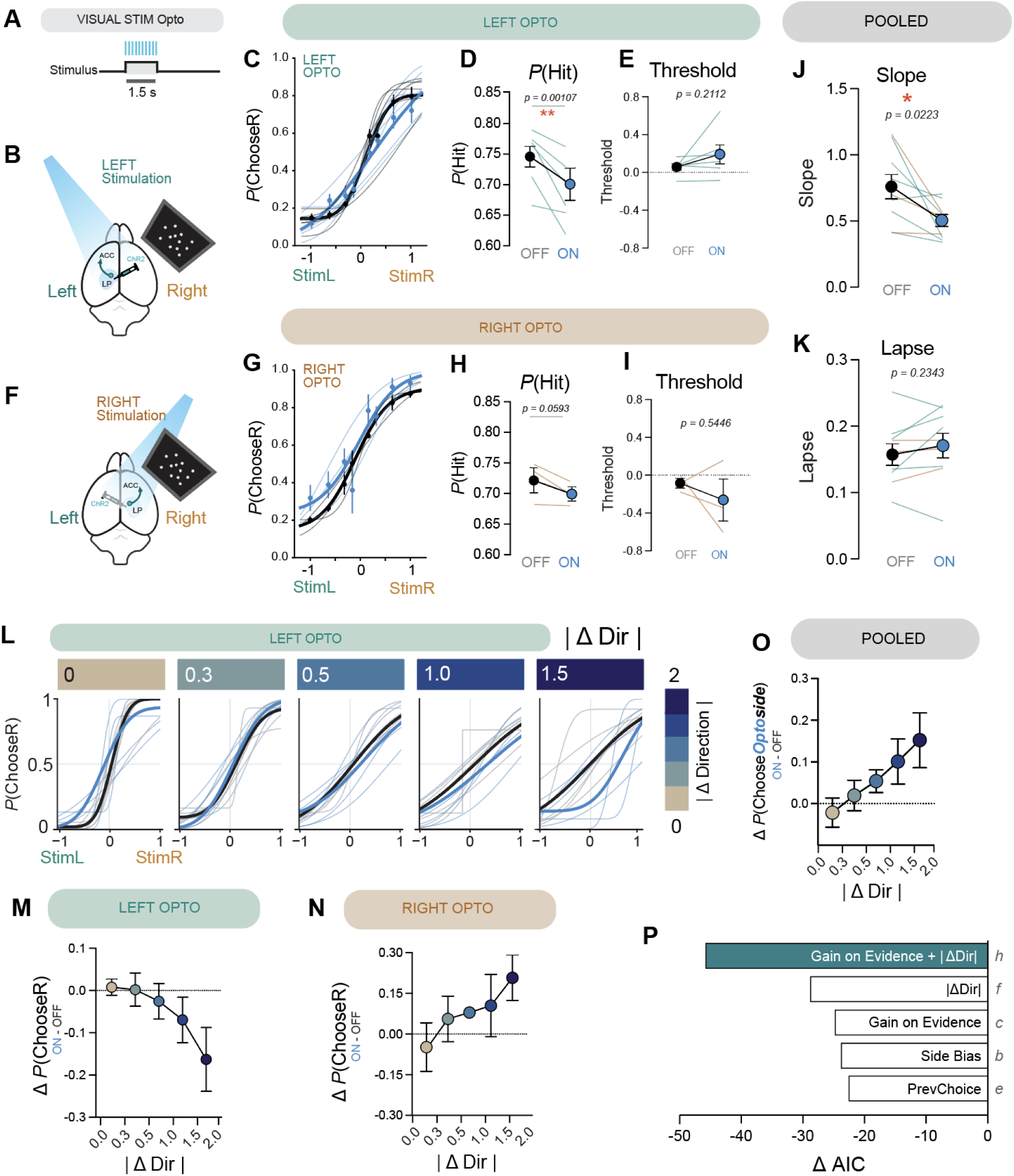
Activating LP-ACC projections biases choice in a |Δ Dir |-dependent manner. (**A**) Trial timeline showing visual stimulus presentation (1.5 s) and optogenetic stimulation delivered during the stimulus period in a random 25-30% of trials. (**B**) Schematic of left LP axons expressing ChR2 stimulated over the ACC. (**C**) Psychometric functions (P(ChooseR)) for left LP-ACC opto-stimulation trials (ON; blue) and interleaved control trials (OFF; black). Thin lines indicate individual animals; thick lines indicate population mean. (**D**) Hit rate (P(Hit)) and (**E**) Psychometric threshold upon left LP-ACC stimulation. (**F** to **I**) Same as B-E but for optogenetic stimulation of right LP axons over ACC. (**J**) Psychometric slope and (**K**) Lapse rate upon unilateral LP-ACC stimulation, pooled across stimulation sides. (**L**) Psychometric functions for left LP-ACC stimulation trials, stratified by |ΔDir| (absolute change in stimulus direction from the previous trial). (M and N) ΔP(ChooseR) as a function of |ΔDir| for (**M**) left and (**N**) right LP-ACC stimulation. (**O**) Change in probability of choosing the optogenetically stimulated side (ΔP(ChooseOptoside) as a function of |ΔDir|, pooled across stimulation sides. (**P**) Model comparison by ΔAIC relative to a baseline model without any Opto term (Table 4, model a). Models are ordered by best fit on top. Models incorporating optogenetic modulation of evidence gain (model c) or |ΔDir|-dependent bias (model f) improved fit relative to static side bias (model b) or previous-choice-dependent modulation (model e). The best-fitting model (model h) is highlighted and combined global gain modulation of sensory evidence (DirCoh:Opto) with side-specific |ΔDir| bias (AbsDiff:OptoSide). Full likelihood ratio statistics and model comparisons are reported in Table 4.

Unilateral stimulation of left LP-ACC axons led to an overall impairment in hit rates (Fig. 2C and D), without a corresponding shift decision threshold (Fig 2E), arguing against disruption by simple motor biases. Surprisingly, stimulation of right LP-ACC axons – ipsilateral to stimulus presentation – also impaired performance (Fig. 2G and H), despite the lack of direct contralateral visual input. As with left stimulation, decision thresholds remained unchanged (Fig 2I). Optogenetic activation of either hemisphere significantly reduced psychometric slopes (Fig. 2J), indicating a selective reduction in sensitivity and influence of sensory evidence on choice formation, rather than a fixed bias. In contrast, both lapse rates (Fig. 2K) and rate of trial omissions (fig. S3B) were unchanged, suggesting that reduced performance is unlikely explained by attentional disengagement or motor errors. Together, these findings indicate that LP-ACC axons contribute to evidence integration to guide decisions, rather than relaying contralateral visual input.

We next asked whether the reduction in psychometric slope following LP-ACC perturbation reflected history-dependent modulation of evidence sensitivity. When we stratified trials by |ΔDir|, optogenetic effects became progressively greater with increasing contrast between current and previous sensory evidence. Specifically, during left LP-ACC stimulation, mice became less likely to make rightward choices (reduced P(ChooseR)) as |ΔDir| increased (Fig. 2L and M, fig S3F-H). In contrast, stimulating the right LP-ACC axons produced mirror symmetric effects – with an increased likelihood of rightward choices with greater |ΔDir| (Fig. 2N, fig. S3I-K). When pooled across stimulation hemispheres, unilateral optogenetic stimulation of LP-ACC axons increasingly biased choices towards action associated with the side of optogenetic stimulation (ipsilateral) as |ΔDir| increased (Fig. 2O). This scaling suggested that stimulation interacts with trial-to-trial sensory differences rather than producing a fixed motor bias.

Consistent with this, models incorporating a |ΔDir|-dependent, hemisphere-specific bias (|ΔDir|:OptoSide; model f) provided a better account of the data than models based on fixed side bias (model b) or previous choice modulation (model e; Fig. 2P, Table S4). However, optogenetic stimulation also produced a global reduction in psychometric slope (Fig. 2J), raising the possibility that stimulation alters sensitivity to current sensory evidence in addition to generating a |ΔDir|-dependent bias. To test whether these effects reflect separable mechanisms, we compared models including a global evidence interaction (DirCoh:Opto), a |ΔDir|-dependent lateralized bias (|ΔDir|:OptoSide), or both terms simultaneously. Each interaction independently improved model fit, and the best-fitting model required both DirCoh:Opto and |ΔDir|:OptoSide terms (model h, Table S4). Thus, LP-ACC stimulates operates through two separable components – a |ΔDir|-dependent, hemisphere-specific bias and a change in evidence weighting. Importantly, the direction of this effect depended on the stimulated hemisphere and scaled with |ΔDir|, revealing a lateralized mechanism by which sensory history governs decision updating.

Whereas unilateral LP-ACC perturbation produced detrimental lateralized effects that scaled with the difference between successive stimuli, bilateral stimulation abolished this lateralization and did not impair performance (fig. S4). This contrast is consistent with unilateral optogenetic effects arising from asymmetric disruption of a normally bilateral LP-ACC input.

Disruption of |ΔDir|-dependent decision-making was specific to perturbing LP-ACC inputs. Illumination at 593nm, which does not activate ChR2, produced no behavioral effects (fig. S5), arguing against non-specific light artifacts. Furthermore, optogenetically stimulating left LP axons in the primary visual cortex (V1) also did not affect behavior or reproduce |ΔDir|-dependent choice biases (fig. S6), suggesting that the observed effects did not reflect indirect activation of other cortices or non-specific disruption of visual processing.

To test whether LP-ACC perturbation caused persistent effects beyond the stimulation period, we examined trials following optogenetically stimulation. Behavioral performance recovered after optogenetic trials regardless of whether the optogenetic trial led to a hit or an error (fig. S7). In addition, optogenetic activation during reinforcement epoch similarly had no effect on choices in the next trial (fig. S8). Both findings indicate that LP-ACC perturbation affected behavior transiently during evidence evaluation and did not disrupt memory of prior stimuli or outcomes. Taken together, these optogenetic results show that LP-ACC activity biases decisions in a |ΔDir|-dependent manner by selectively shaping how current sensory evidence is evaluated based on recent sensory context.

### LP-ACC axons represent visual stimuli with task-gated low-dimensional geometry

The optogenetic results suggest that LP-ACC inputs causally influence how sensory evidence guides choice in light of recent sensory history, rather than providing a simple feedforward visual signal. To determine the nature of information provided by LP, we performed two-photon imaging of LP-ACC axons during task performance and passive viewing (Fig. 3A, fig. S9). During task engagement, LP-ACC axons exhibited robust, stimulus-evoked responses (Fig. 3B, fig. S11A-B) that collectively discriminated stimuli defined by direction and coherence, whereas this stimulus information was much weaker under passive viewing in naive mice (Fig. 3C). This task dependence suggests that LP-ACC activity reflects behaviorally relevant sensory variables rather than bottom-up visual responses alone.

**Figure 3.**
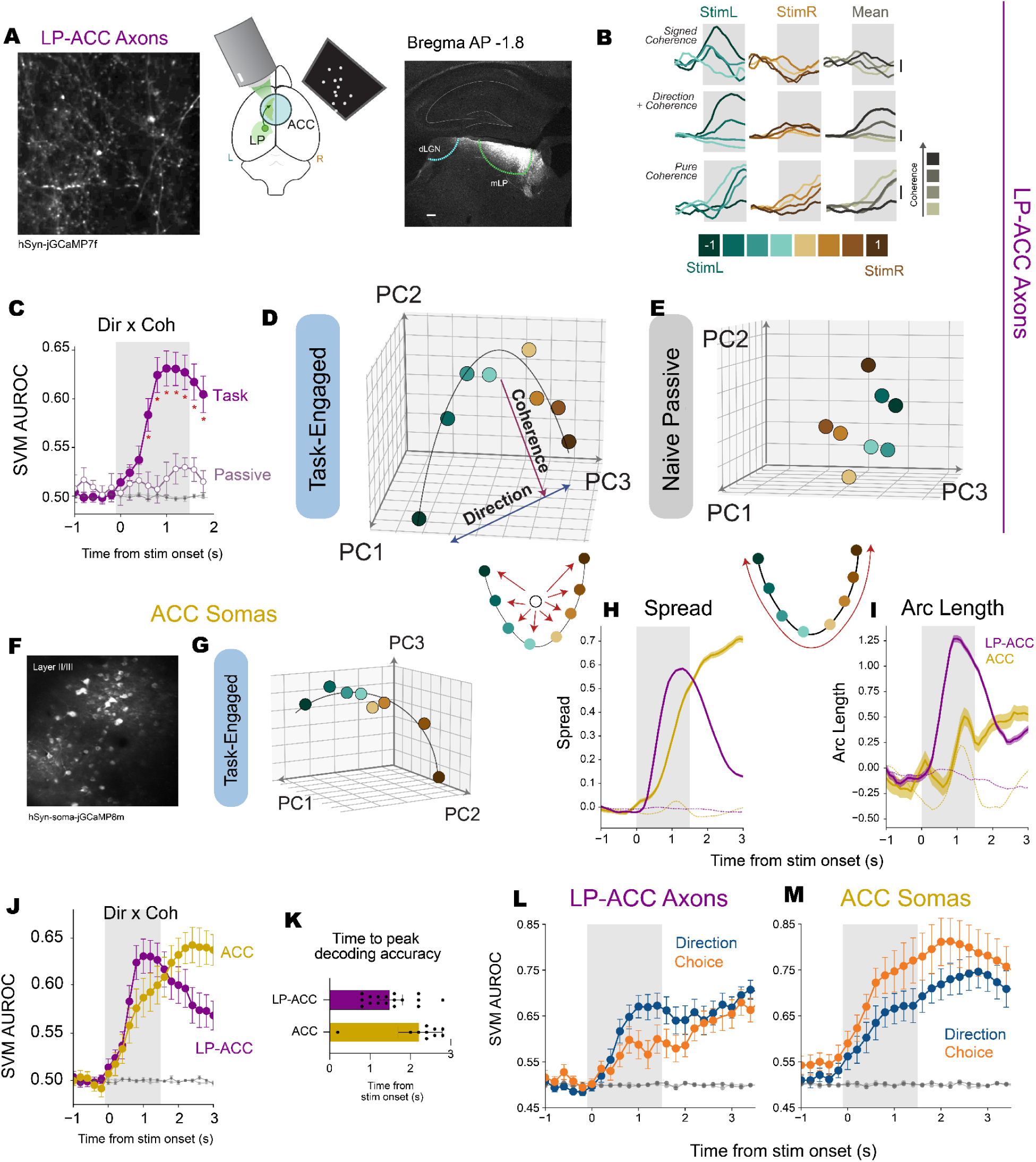
LP-ACC axons represent visual stimuli along a curved low-dimensional manifold. (**A**) Schematic of two-photon calcium imaging of LP axons in ACC during task performance, and example field of view showing GCaMP-expressing LP axons in ACC (left) and representative confocal micrograph showing medial LP targeting (right). (**B**) Example trial-averaged traces of single LP-ACC axons with activity that scales with signed coherence (top), with preferential activity or left stimuli but also scales monotonically with stimulus coherence (middle), and representing pure coherence encoding without direction discriminability (bottom). (**C**) Cross-validated SVM decoder performance (AUROC) trained to discriminate each of the 8 presented stimuli (2 directions x 4 coherences) perform much better in the task-engaged condition compared to in naive mice passively viewing identical stimuli. Red asterisks indicate significant difference (p < 0.05) in decoding performance at that time-point, using two-tailed t-test. (**D**) LP-ACC population activity projected onto the first 3 PCs for unique presented stimuli when mice were engaged in the task. The representation organizes into a curved manifold that simultaneously represents stimulus direction and coherence. (**E**) Same as D but in naive mice passively viewing identical stimuli; LP-ACC population does not have apparent curved manifold organization. (**F**) Schematic of two-photon calcium imaging of ACC during task performance, and example field of view showing GCaMP-expressing LP axons in ACC. (**G**) ACC population activity projected onto first 3 PCs during task engagement. (**H** and **I**) Manifold metrics, pre-stimulus baseline-subtracted. Spread (H), representing dispersion of stimulus representations from centroid, and Arc length (I), which measures path distance through ordered signed coherence, for LP-ACC (purple) and ACC (yellow). Shading indicates SEM across iteration. (**J**) Cross-validated SVM decoder performance (AUROC) like in (C) but comparing with ACC (yellow) populations. (**K**) Comparison of time to peak decoding accuracy for Dir x Coh decoders in J. Each dot represents a single imaging session. (**L** and **M**) Cross-validated SVM decoder performance (AUROC) decoding binary direction (blue) or choice (orange) for (L) LP-ACC axons and (M) ACC somas. For task-engaged LP-ACC decoding (C, J and L), n =17 sessions, 5 mice. For passive LP-ACC decoding (C), n = 13 sessions, 5 mice. For task-engaged ACC decoding (J and M), n = 9 sessions, 3 mice.

While decoding demonstrates that LP-ACC activity distinguishes stimulus direction, the heterogeneous tuning for direction and coherence across individual LP axons (Fig. 3B, fig. S11) raised the question of how graded evidence for stimulus could be represented across the LP-ACC population. To address this, we summarized trial-averaged population activity for each stimulus using cross-validated principal component analysis (PCA; first 3 principal components: 34.7% total explained variance). During task performance, LP-ACC population responses varied smoothly and in an orderly manner with graded changes in stimulus direction and strength, giving rise to a curved trajectory aligned with signed stimulus coherence (Fig 3D). We refer to this continuous curved trajectory as a low-dimensional curved manifold. Along this manifold, population activity jointly reflected stimulus direction and stimulus difficulty, rather than clustering into discrete stimulus categories (fig. S10A). Notably, this low-dimensional geometry in LP-ACC activity was not evident in naive mice during passive viewing despite identical sensory input (Fig. 3E). Moreover, the same structure was observed on omission trials lacking overt licking behavior (fig. S10), ruling out a movement-related origin. Together, these results demonstrate that this organization in graded sensory evidence in LP-ACC population geometry is present during task performance but not during passive viewing.

As a point of comparison to established cortical decision signals, we examined population activity in ACC neurons during the same task. Consistent with prior work, ACC populations exhibited low-dimensional structure during the task (Fig. 3F-G, fig. S10F-G and J-K) but the timing and alignment of this structure differed markedly from LP-ACC inputs. During stimulus evaluation, LP-ACC axons exhibited a pronounced expansion of population geometry, reflected by larger spread and longer arc length (Fig. 3H-I). Increased spread indicates greater separation of stimulus representations, whereas longer arc length reflects more extensive and orderly monotonic organization of graded sensory evidence along the curved manifold (Materials and methods).

Spread was consistently larger in LP-ACC populations than ACC during the stimulus period, whereas ACC spread increased later to exceed LP-ACC at its peak (Fig. 3H). In contrast, arc length remained larger in LP-ACC throughout the stimulus epoch, while ACC exhibited more modest increase later (Fig. 3I).

These temporal and geometric differences were also reflected in decoding analyses. Decoding of graded sensory evidence peaked during stimulus presentation for LP-ACC axons, whereas ACC decoding peaked later near the response (Fig. 3J to K). Moreover, direction decoding dominated in LP-ACC axons during stimulus evaluation, while ACC populations showed stronger choice than direction decoding during the response period (Fig. 3L-M, fig. S10L-M), highlighting a temporal shift from sensory-aligned to choice-aligned population representations across the LP to ACC pathway.

### Sensory history reshapes LP-ACC population geometry during stimulus evaluation

Having established that LP-ACC population activity is preferentially aligned with sensory variables during evidence evaluation, we next asked how recent sensory history modulates representations. A prerequisite for such modulation is that information about the previous stimulus is available during the period when current sensory evidence is evaluated. Consistent with this, previous stimulus direction could be reliably decoded from LP-ACC population activity during the inter-trial interval and early in the subsequent trial (Fig. 4A). Previous stimulus direction decoding generalized across trial outcomes – even when the animal’s choice was incorrect – indicating dominance of sensory rather than choice history (fig. S12A). Decoding accuracy declined during stimulus presentation, consistent with a transition from history-dominated to stimulus-dominated representations.

**Figure 4.**
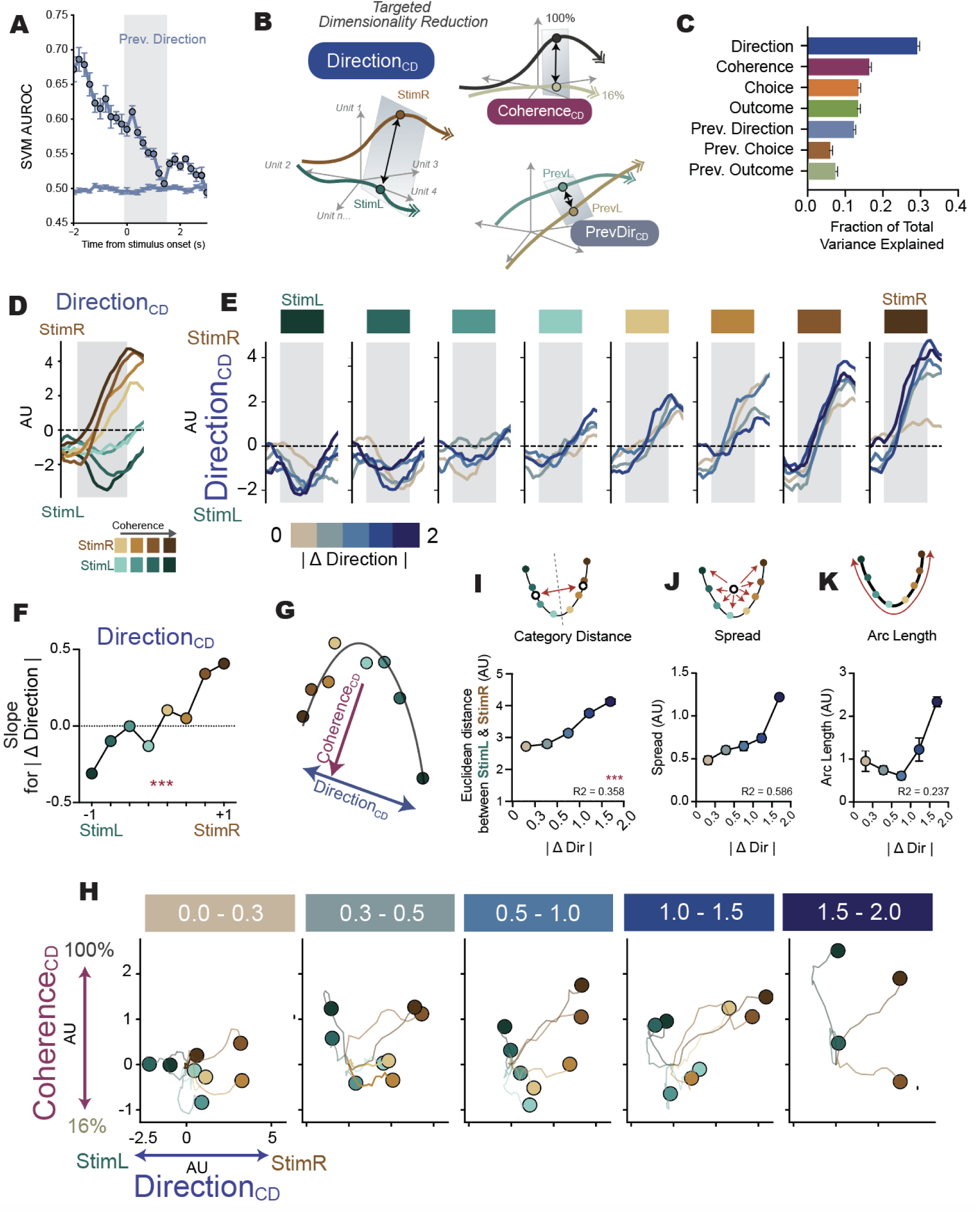
Sensory history reshapes LP-ACC population geometry during stimulus evaluation. (**A**) Cross-validated SVM decoder performance (AUROC) decoding of previous direction (blue). (**B**) Schematic of targeted dimensionality reduction (TDR) analysis used to define task-relevant coding dimensions (CDs) such as Direction, Coherence and Previous Direction from LP–ACC population activity. (**C**) Fraction of total variance explained by each task CD in LP-ACC axonal populations. (**D**) Projection of LP-ACC population activity onto the Direction_CD_ (arbitrary units, AU) show graded representation of stimulus signed coherence. (**E**) Trial-averaged projections onto the Direction_CD_ (AU) for each of the 8 unique stimuli, separated by |ΔDir| (absolute change in stimulus direction from the previous trial), showing systematic shifts with increasing |ΔDir|. (**F**) Quantification of projection shifts along Direction_CD_ as a function of |ΔDir| for left- and right-associated stimuli. Modulation as quantified by slopes for |ΔDir| for each stimulus. Slope derived from least squares fit for |ΔDir| to mean projection (AU). One-way repeated measures ANOVA on effect of stimulus direction on slopes (p = 1.65 x 10-43, F(7,525) = 37.9, ηp2 = 0.336, over 75 iterations). (**G**) Schematic illustrating the Direction_CD_ and Coherence_CD_ from TDR that recapitulates curved manifold observed in PCA. (**H**) Population trajectories projected into the joint Direction_CD_-Coherence_CD_ subspace (AU), revealing a curved manifold whose structure expands with |ΔDir|. Large dots represent each trajectory at the end of the stimulus evaluation period (1.5s after stimulus onset), lines represent their trajectory from baseline (subtracted) upon stimulus onset. (**I** to **K**) Manifold Metrics. (I) Category distance, measured as euclidean distance between mean StimL and StimR representations, increases monotonically as a function of |ΔDir|. (J) Spread, representing dispersion of stimulus representations from centroid, also increases as a function of |ΔDir|. (K) Arc length, path distance through ordered signed coherence, has a U-shaped relationship as a function of |ΔDir|.

The persistence of previous sensory information into the early stimulus period indicates that LP-ACC populations concurrently represent recent and current sensory evidence. Encoding models further revealed widespread multiplexing of current and previous trial variables in single axons that could not be explained by motor correlates alone (fig. S13), motivating a population-level analysis of how task variables are integrated within LP-ACC representations.

Behavioral and optogenetic results identified trial-to-trial differences in sensory evidence (|ΔDir|) as a key determinant of choice. We therefore asked whether LP-ACC representations of current sensory evidence are systematically reorganized as a function of |ΔDir|, and whether this modulation acts along the same dimensions that encode sensory evidence itself. To address this, we used targeted dimensionality reduction (TDR), a method that identifies the population activity axes that best explains variance related to a specific task variable (Fig. 4B). We can then isolate population coding dimensions (CDs) to quantify how activity along these dimensions varies with |ΔDir|. Current stimulus direction explained the largest fraction of LP-ACC population task-related variance, while PrevDirectionCD explained greater total variance than PrevChoiceCD (Fig. 4C), reinforcing that LP-ACC history representation is sensory-, rather than choice-dominant.

We first examined activity along the coding dimension corresponding to stimulus direction (Direction_CD_). Projections onto Direction_CD_ revealed responses to the same visual stimulus were systematically shifted depending on the magnitude of change in stimulus direction relative to the previous trial (|ΔDir|; Fig. 4D-F). These shifts occurred along the same coding dimension used to represent current sensory evidence, rather than along a separate history-specific or orthogonal axis.

We next asked how this axis-level modulation shapes the global geometry of sensory representations. In the joint Direction_CD_ and Coherence_CD_ subspace, LP-ACC population activity formed a curved manifold whose structure depended on |ΔDir| (Fig. 4G). When the stimulus direction was similar to the previous (low |ΔDir|), representations were more compact, whereas increasing |ΔDir| was associated with progressive expansion of the manifold (Fig. 4H). Consistent with this expansion, the Euclidean distance between left- and right-associated stimulus representations increased monotonically with |ΔDir| (Fig. 4I), indicating enhanced representational separation when current sensory input deviated more strongly from recent history. Beyond category separation, representations also spanned a larger extent of the manifold with increasing |ΔDir|, as reflected by increases in trajectory spread and arc length (Fig. 4J,K).

Notably, a comparable history-dependent expansion of representational geometry was not observed in ACC neural populations analyzed in the same manner (fig. S14), suggesting that LP-ACC axons provide a history-referenced input that is transformed, rather than directly preserved, in ACC population activity. Together, these results show that trial-to-trial change in sensory evidence is encoded as a graded reorganization of LP-ACC population geometry, preferentially expanding representational space for larger deviations from recent sensory context.

### Contrastive coding of stimulus history in LP-ACC prioritizes trial-to-trial change

The history-dependent expansion of LP-ACC population geometry raises a key question: what coding structure gives rise to a history-dependent rescaling of sensory representations, rather than a uniform shift? To address this, we examined the TDR-defined population coding weights to determine how current and previous sensory evidence are combined across LP-ACC axons.

Population weights defining the Direction_CD_ and PrevDirection_CD_ axes were systematically anti-correlated (Fig. 5A) indicating opponent contributions of current and recent sensory evidence at the population level. This organization predicts that sensory history will rescale stimulus-evoked responses rather than introduce a uniform offset. To illustrate how this population-level opponency is reflected in axonal responses, we examined population-averaged activity of StimR-selective LP-ACC axons grouped by aligned versus contrastive coding weights to the same stimulus. Axons with aligned weights exhibited similar response polarity for current and previous stimulus direction, consistent with a shift-like influence of history (Fig. 5B and C). In contrast, axons with contrastive weights showed opposing modulation, such that responses become suppressed when the previous stimulus direction matched the current stimulus (Fig. 5D and E). The opponent structure was specific to sensory history: Direction and Choice weights were positively aligned (fig. S15A), whereas coupling between Direction and PrevChoice weights was weak and inconsistent (fig. S15B and E), arguing against a generic correlation across task variables.

**Figure 5:**
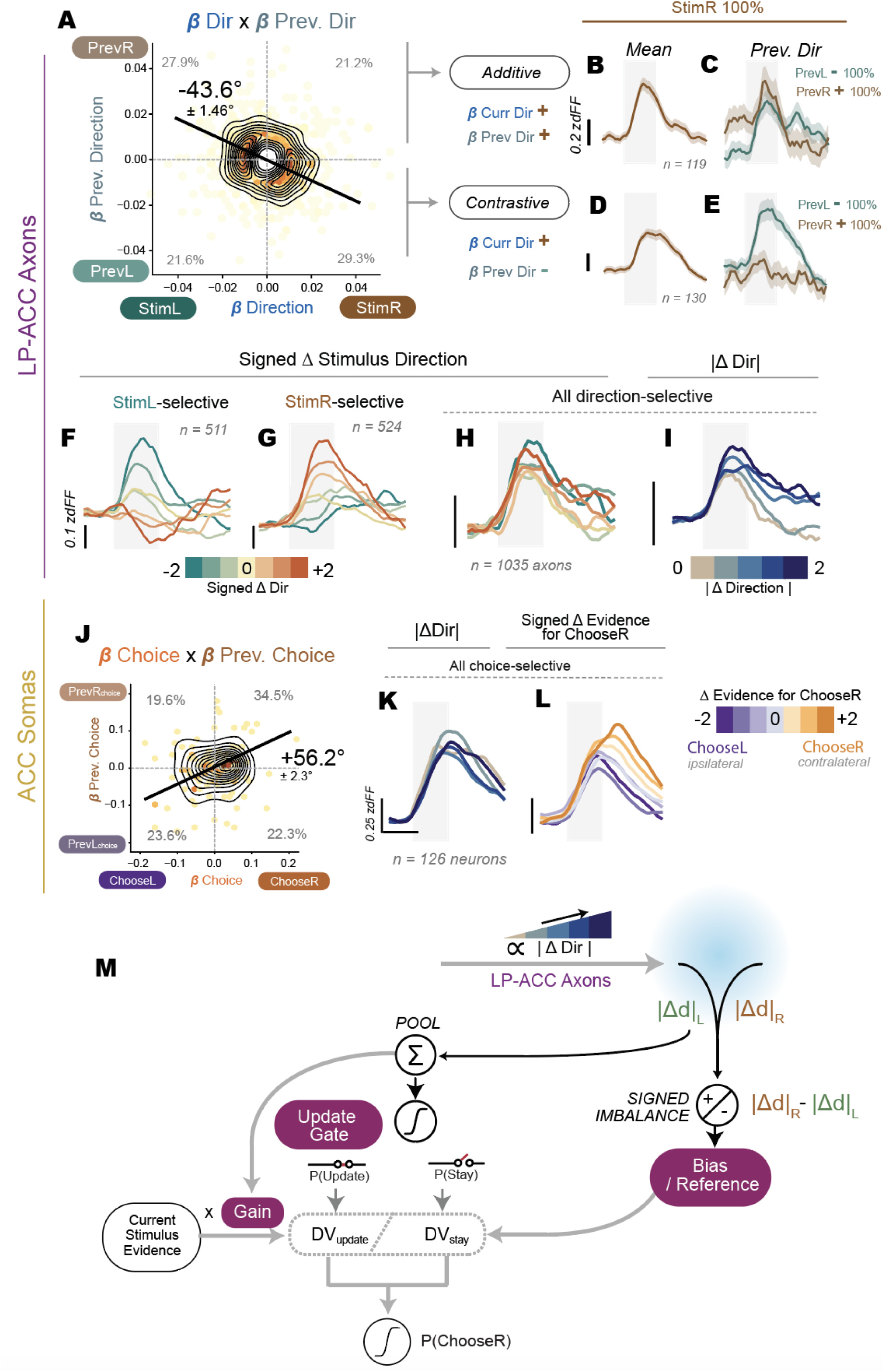
Contrastive sensory history coding in LP-ACC supports history-dependent decision updating. (**A**) Correlation between LP-ACC axons’ current stimulus direction (β Dir) and previous stimulus direction (β Prev.Dir) coefficients demonstrating a predominantly negative relationship indicative opponent encoding. Only top 50% of weights shown in plot for visualization purposes, all analyses were performed on the complete set of weights. Darker colors indicate higher density of units. Percentages indicated the proportion out of all axons that fell in each quadrant. Mean major axis angle reported with SEM across 100 iterations of TDR. (**B** and **C**) Subpopulation of LP-ACC axons with additive aligned weights, selective for both StimR (β Dir +) and PrevR (β Prev.Dir +), averaged across all trials of the same StimR 100% stimulus (B) or stratified by previous direction at 100% coherence (C). As weights are aligned and additive, PrevR responses to the same consecutive stimulus were greater. (**D** and **E**) Subpopulation of LP-ACC axons with contrastive weights, selective for both StimR (β Dir +) and PrevL (β Prev.Dir -), averaged across all trials of the same StimR 100% stimulus (D) or stratified by previous direction at 100% coherence (E). The contrastive weights led to reduced responses when stimulus was repeated (both current StimR and PrevR 100%) while responses were greatest at maximal difference (StimR & PrevL at 100%) from history. B to E scale bars all correspond to 0.2 z-scored dFF. (**F** and **G**) Subpopulation average of StimL- (F) and StimR-selective (G) LP-ACC axons, stratified by binned signed ΔDir (Current Signed Coherence - Previous Signed Coherence), showing that signed ΔDir information is present, relative to axons’ preferred tuning. (**H**) Pooled direction-selective axons no longer show monotonic modulation by signed ΔDir at the level of bulk responses. (**I**) Unsigned |ΔDir| is reflected in bulk responses of pooled direction-selective axons. F to I scale bars all correspond to 0.1 z-scored dFF. (**J**) Correlation between ACC neurons axons’ current choice (β Choice) and previous choice (β Prev.Choice) coefficients showing strong positive relationship, indicating dominantly additive coding of choice history. Other plot details as described for (A). (**K**) Subpopulation average of all choice-selective ACC neurons do no show monotonic scaling by unsigned |ΔDir|. (**L**) Choice-selective axons are modulated by Δ Evidence for contralateral choice (ChooseR). This modulation is signed with strong asymmetric anchoring to contralateral choice. K and L scale bars correspond to 0.25 z-scored dFF. (**M**) Conceptual model linking LP-ACC comparison signals to history-dependent decision updating. LP-ACC axons encode trial-to-trial differences in sensory evidence, indexed by either signed ΔDir or unsigned |ΔDir|. This signal is provided independently to ACC in each hemisphere, with |dL| and |dR| representing signals to each hemisphere which are usually identical under normative conditions. These signals contribute to decision-making through three separable computations. First, the pooled magnitude of LP-derived comparison signals governs the probability of entering an update versus stay regime (“Update Gate”). Second, within the update regime, LP-derived signals modulate the “gain” applied to current sensory evidence, scaling the influence of incoming evidence on the decision variable. Third, a signed “bias/reference signal” is generated in ACC due to interhemispheric competitive interactions (|Δd|_R_ - |Δd|_L_) if LP-ACC signals are asymmetric. Choice probability (P(ChooseR)) is determined by distinct decision variables computations in either the stay or update regime. Optogenetic stimulation is simulated by perturbing the |Δd|_R_ and/or |Δd|_L_ signals upstream (indicated by blue illuminated area). This model schematic depicts the unsigned comparator variant tested – other variants are described in Supplementary Note 1. This framework accounts for |ΔDir|-dependent switching, lateralized effects of unilateral LP-ACC perturbation, and cancellation under bilateral stimulation.

When responses were grouped by stimulus tuning, both StimL- and StimR-selective axons showed systematic scaling with signed stimulus difference relative to their preferred direction, with no systematic bias for one stimulus direction over the other (Fig. 5F and G). When activity was pooled across axons, signed stimulus-difference information was not prominently expressed in bulk averages (Fig. 5H), despite remaining linearly separable in population space, as revealed by low-dimensional population geometry (Fig. 4). In contrast, pooling emphasized the magnitude of trial-to-trial sensory change (|ΔDir|; Fig 5I), which we previously observed to be associated with an expansion of the population manifold and increased separation between stimulus representations (Fig. 4). Together, these analyses show that LP-ACC population activity jointly conveys stimulus identity and sensory change magnitude: stimulus identity is embedded in the geometry of population responses, while |ΔDir| is reflected both through geometric expansion and aggregate population activity.

We next turned to ACC, where population activity is dominated by choice-related variance (fig 15H). We therefore examined the relationship between current and previous choice representations. In contrast to the opponent organization in LP-ACC, ACC neurons showed predominantly aligned, positively correlated coding for current and previous choice (Fig. 5J, fig S15I to M), consistent with reinforcement of action history and a shift-like bias in choice representations rather than contrastive integration. Accordingly, unlike LP-ACC, monotonic scaling with |ΔDir| was no longer apparent in ACC bulk responses (Fig. 5K). Instead, ACC bulk activity scaled monotonically with the signed difference in choice-evidence signal, driven by aligned contributions of current and previous choice, as well as the dominance of contralateral (ChooseR) choice-selective neurons (Fig. 5L, fig. S15N to P). In sum, these imaging results delineate a transformation from contrastive, stimulus-centered representations in LP-ACC to aligned, choice-centered representations in ACC.

### A model linking LP comparison signals to |ΔDir|-dependent choice updating

Integrating the imaging and optogenetic results, we therefore implemented a reduced model in which LP provides stimulus-anchored comparison signals independently to each hemisphere (Fig. 5M, Supplementary Note 1-2); the information sent to each hemisphere is assumed to be identical. In the model, LP-derived comparison signals play two separable functions: their pooled magnitude controls entry into an update regime (“Update Gate”), while their interhemispheric difference provides a signed imbalance that biases choice direction upon updating (“Bias/Reference”). As |ΔDir| information sent to each hemisphere is assumed to be identical, the net bias is always zero except when simulating unilateral optogenetic stimulation.

Here, behavior defaults to a win-stay strategy unless |ΔDir| information provides sufficient drive to enter the update regime – an internal cognitive state, rather than an action state, and thus does not necessarily involve choice switching. Within the update regime, the model incorporates increasing |ΔDir|-dependent gain on sensory evidence (“Gain”), consistent with the expansion of stimulus representations observed in LP-ACC population activity. This allows strong and clear differences in evidence to overcome the intrinsic cost of switching and restore performance, and was a mechanism required to reproduce the non-monotonic psychometric sensitivity observed in normative behavior (Fig. 1I to L, fig. S16A to F). Thus, behavior emerges from a mixture of persistence (“stay”) and change-responsive (“update”) regimes, in which LP-derived |ΔDir| signals provide information that guides updating without directly encoding choice.

Simulating optogenetic modulation as changes in LP-derived ΔDir comparison signals applied prior to ACC readout, the model reproduced |ΔDir|-dependent entry into update states, lateralized biases under unilateral perturbation, and cancellation under bilateral stimulation (fig. S16G to Y). Crucially, the model reproduced the observed |ΔDir|-dependent scaling of optogenetic effects, as larger |ΔDir| increased both update probability and the magnitude of the induced bias through greater interhemispheric mismatch. Perturbing LP-derived comparison signals prior to ACC readout was sufficient to account for these effects across various perturbation schemes (fig. S16G to Y, Supplementary Note 1). Notably, selective modulation of the bias term alone was sufficient to reproduce the primary lateralized optogenetic phenotype (Supplementary Note 1, fig. S17B). However, reproducing the full baseline behavioral structure – including the non-monotonic psychometric sensitivity (Fig. 1K-L) – required the mixture of stay and update regimes that governs |ΔDir|-dependent gain (Supplementary Note 1, fig. S17). Together, these simulations indicate that LP-derived |ΔDir| signals inform when updating occurs and bias its direction, without directly encoding choice.

## Discussion

Perceptual decision-making involves comparing current sensory evidence to recent experience, yet how such comparisons are implemented at the circuit level remains unclear. Here, we identify a thalamocortical mechanism in which this comparison is realized via a contrastive population code in LP projections to ACC. LP-ACC axons encode current sensory evidence relative to the previous trial via an opponent coding structure, rather than a uniform shift. As a result, population activity is organized to emphasize changes from recent sensory history: low-dimensional population manifolds representing stimuli expand with increasing trial-to-trial changes in evidence, embedding information about stimulus identity and change in sensory evidence within population structure. By emphasizing deviations from recent history, this contrastive population code is well suited to influence when incoming evidence leads to an update.

At a mechanistic level, our results support a role for LP-ACC inputs in reshaping the population representation of sensory evidence through comparison-level gain. Contrastive coding of current relative to recent sensory evidence gives rise to an expansion of population geometry with increasing differences between successive stimuli, reparameterizing decision-relevant evidence. Critically, this rescaling need not reflect changes in low-level sensory encoding: rather than globally enhancing or suppressing sensory evidence, LP-ACC signals bias how evidence is represented and compared, and may modulate the effective gain of these representations in a history-dependent manner.

Consistent with a comparison-level role, our findings argue against a model in which LP-ACC projections act as a feedforward visual pathway. Unilateral optogenetic perturbation produced mirror-symmetric behavioral effects, including when stimulation was applied to the hemisphere lacking direct sensory input (Fig. 2). A strictly feedforward sensory pathway would instead predict asymmetric effects tied to sensory drive. This view is further supported by task-gated visual representations (Fig. 3) and the presence of history-dependent modulation (Fig. 4 and 5), demonstrating that LP-ACC representations are shaped by behavioral context. Consistent with this interpretation, medial pulvinar circuits projecting to frontal cortex are relatively less constrained by visual topography and instead enriched for associative, multisensory and motor-related inputs (*17,23,35*), compared to pulvinar subdivisions embedded within early visual cortical loops (*11,12,18–22,36–40*).

This circuit-level interpretation also clarifies the temporal locus of LP-ACC influence. History-dependent biases had been described as shifts in initial conditions or bias terms in evidence accumulation models (*41*), and have been linked to distributed cortical representations of prior information (*42*). Several cortical regions, most notably posterior parietal cortex (PPC), have been implicated in mediating such biases, with inactivations during the ITI reducing history-dependent influences on subsequent choices (*43,44*). In contrast, despite robust representation of previous-trial information in LP-ACC axons during the ITI, ChR2 activation of LP-ACC during this epoch had little impact on behavior (fig. S8). Instead, LP input to ACC exerted its primary influence during evaluation of the current sensory evidence, biasing its representation relative to recent history. This temporal dissociation suggests that LP-ACC contributes to history-dependent decision biases through modulation of ongoing evidence evaluation, rather than by setting a persistent bias state prior to stimulus onset.

Consistent with a role during stimulus evaluation, our results indicate that LP-ACC inputs regulate when and how sensory evidence is incorporated to drive updating, rather than simply biasing decisions toward or away from recent choices. In both imaging and modeling, contrastive comparison signals scaled with the magnitude of differences in sensory evidence and gated entry into an update regime, linking trial-to-trial comparison to changes in evidence evaluation that ultimately shape action. This comparison-based update away from a default win-stay policy provides a concrete circuit mechanism for context-dependent control of decision updating and is consistent with theoretical frameworks that emphasize predictive and context-dependent modulation of cortical processing by the pulvinar (*12,13,36–38*). We formalized behavior as a mixture of stay and update regimes, with LP-derived |ΔDir| signals particularly important for supporting updating (Fig. 5M, Supplementary Note 1). Nevertheless, whether such history-dependent modulation of evidence leads to behaviorally optimal choices across task contexts, or instead reflects circuit-level constraints or priorities independent of task demands, remains an open question. Our results instead support a more constrained interpretation in which decisions are shaped by a difference-based evaluation of current sensory evidence relative to recent sensory history. Under this view, LP inputs contribute information relevant for determining when sensory evidence is sufficiently different to support updating a prior choice, but this computation is accompanied by a switch-cost. We therefore emphasize that the observed history dependence reflects a mechanistic property of evidence evaluation rather than a flexible strategy designed to optimize behavioral performance.

At the population level, history-dependent modulation of sensory evidence is implemented in LP-ACC through scaling of representational geometry and transformed downstream within ACC. We showed that both LP-ACC axons and ACC neurons represent the sensory evidence in a structured curved geometry (Fig. 3, fig. S10), extending low-dimensional manifold representations of decision variables beyond cortical circuits (*45,46*) to thalamic structures embedded in recurrent decision networks. Importantly, we observed expansion of geometric representations that scaled with the difference in sensory evidence across trials in LP-ACC axonal populations (Fig. 4 and 5), but not in ACC (fig. S14), where history induces more additive, shift-like biases along choice-centered dimensions. This dissociation suggests that a common low-dimensional scaffold for evidence representation can be differentially modulated in a region-specific manner, reflecting distinct computational roles within the decision network and respective tuning population tuning (*47*).

LP-ACC input emphasizes contrastive, history-relative sensory evidence, while ACC transforms these inputs into choice-aligned representations through competitive integration. Critically, choice representations in ACC are lateralized and evaluated through interhemispheric competition, such that balanced activity across hemispheres is required for unbiased choice (*48–50*). Because LP-ACC inputs are anchored to sensory variables rather than choice, and also did not show systematic preference for one side, their influence on behavior must therefore be mediated through how they perturb this competitive ACC readout. Consistent with this framework, we found that unilateral – but not bilateral – perturbation of LP input led to biased choice, mirroring effects of lateralized perturbations previously reported in ACC. The abolition of bias under bilateral activation indicates that optogenetic effects depend on interhemispheric imbalance, and thus engage downstream competitive ACC circuitry.

Our model shows that interhemispheric balance can be disrupted in multiple ways. Distinct perturbations of LP-derived comparison signals – such as gain or noise – were each sufficient to produce lateralized behavioral effects after readout by ACC. Thus, while the optogenetic results implicate LP-derived comparison signals as a critical, sensory-centered input to ACC decision circuitry, the model does not uniquely specify how those signals are altered by stimulation. Rather, behavioral biases emerge through competitive integration as LP-derived signals are transformed within ACC into action-centered representations. How this transformation is implemented at the microcircuit level remains an open question. LP axons preferentially target superficial ACC layers (*24*), where they are well positioned to interact with local inhibitory and recurrent circuitry (*51–56*), providing a plausible substrate through which sensory-centered comparison signals could be converted into action-selective biases.

Viewed in the broader context of pulvinar function, comparison-based rescaling of sensory evidence by LP-ACC inputs aligns with growing evidence that the pulvinar supports predictive and contrastive computations (*7,12,13,57*). Previous work and models have shown that pulvinar-cortical circuits can compute differences between concurrently competing stimuli (*58*), encode confidence-related signals (*57*), or report mismatches between sensory input and self-generated actions such as locomotion or eye movements (*22,36–38*). In contrast, our results provide evidence that LP-ACC projections encode how current sensory evidence deviates from internally maintained, history-dependent expectations, rather than mismatches between simultaneously present sensory or visuomotor signals. This specialization is consistent with the heterogeneous nature of pulvinar outputs, which convey distinct forms of predictive and contextual information to different cortical targets (*10–12, 25, 57*). Accordingly, the LP-ACC pathway illustrates how such history-dependent predictive computations can be routed into prefrontal decision-making circuits.

Prior experience fundamentally shapes how new sensory information is prioritized during decision-making. By encoding current sensory evidence relative to recent history, LP-ACC inputs provide a mechanism for emphasizing deviations from prior experience, thereby shaping when incoming evidence is likely to drive a change in action. Our work positions the pulvinar as an active contributor to perceptual filtering, regulating the influence of sensory signals based on recent history. Such context-dependent modulation may support efficient use of neural processing resources by selectively amplifying behaviorally informative changes in sensory input.

## Acknowledgments

We thank Taylor Johns for lab management and members of the Sur laboratory for many discussions and comments. We also thank Leopoldo Petreanu and former lab members for pointers on beginning the 2AFC RDK discrimination behavior.

## Funding

National Institutes of Health grant R01MH126351 (MS)

National Institutes of Health grant R01MH133066 (MS)

National Institutes of Health grant R01NS130361(MS)

Multidisciplinary University Research Initiatives Grant W911NF2110328 (MS)

The Picower Institute Innovation Fund (MS)

Simons Foundation Autism Research Initiative through the Simons Center for the Social Brain (MS)

A*STAR National Science Scholarship (BS-PhD) (YNL)

## Author contributions

Conceptualization: YNL, MS

Investigation: YNL, AN, AB, CL, SA, MJ, MS

Visualization: YNL

Funding acquisition: MS

Supervision: MS, MJ

Writing – original draft: YNL

Writing – review & editing: YNL, SA, AN, MS, MJ

## Competing interests

Authors declare that they have no competing interests.

## Data and materials availability

All data necessary for evaluating the conclusions are available in the main text or the supplementary materials.

## Supplementary Materials

Materials and Methods

Supplementary Note 1 to 2

Figs. S1 to S17

Tables S1 to S5

## Notes

### Competing Interest Statement

The authors have declared no competing interest.

### Summary of Updates

Added optogenetic experiments, imaging of ACC neurons and theoretical model

