## Supplementary for "Sensory History Shapes Contrastive Neural Geometry in LP/Pulvinar-Prefrontal Cortex Circuits"

**Supplementary Note 1: Behavioral model details**

**Model design:**
The model was designed as a simplified framework to link LP-derived sensory comparison signals to trial-by-trial choice updating. First, a central design feature of the model is that choice formation operates in two regimes with different sensitivity to sensory evidence. |ΔDir| governs gating between these regimes ("Update Gate", Fig. 5M). In the stay regime, which dominates when trial-to-trial sensory differences (|ΔDir|) are small, choices are driven by a high-gain decision variable, resulting in steep psychometric slopes that reflect benefits of choice persistence rather than enhanced sensory discriminability. In contrast, when the model enters an update regime driven by larger |ΔDir|, the effective slope is initially reduced, introducing a transient cost in sensitivity at the moment of updating. However, in this update regime, larger |ΔDir| values make the current sensory evidence more informative about the correct choice, increasing its impact on the decision variable ("Gain", Fig. 5M). This increase in behavioral sensitivity reflects a property of the decision variable rather than an explicit model of sensory discriminability in LP.

Notably, this structure is qualitatively consistent with our imaging data. In LP-ACC axons, overall activity and stimulus discriminability were lowest at small |ΔDir| values and increased with greater |ΔDir|, even though behavioral performance is highest at low |ΔDir| due to choice persistence. Thus, both the model and the data capture a dissociation in which stable behavior at low |ΔDir| does not necessitate strong sensory differentiation, whereas increasing |ΔDir| engages updating and coincides with stronger LP-ACC sensory signals. As an intended consequence of this design, psychometric sensitivity varies non-monotonically with |ΔDir| – with high sensitivity at low |ΔDir| due to choice persistence, reduced sensitivity at intermediate values during updating, and partial recovery at larger |ΔDir| as sensory drive increases (fig. S16A – as we observe in normative behavior (Fig 1I and L). Thus, the P(Update) regime and the gain modulation is essential for reproducing normative |ΔDir|-dependent behavior (fig. S17C).

Second, interhemispheric differences in LP-derived signals are allowed to bias the direction of choice ("Bias/Reference", Fig. 5M). Note that interhemispheric differences are usually equal under normative behavior and it is only under asymmetric stimulation conditions (such as unilateral optogenetic stimulation) that this term is non-zero. This design choice was motivated by the observation that LP responses are anchored to sensory variables rather than strongly biased toward a particular choice. Accordingly, the model does not assume that LP encodes choice direction directly, but instead tests how |ΔDir| signals, whether preserved as signed differences or reduced to unsigned magnitudes, can give rise to lateralized choice biases when read out through an interhemispheric opponent structure and incorporated into the decision variable.

In summary, the model specifies three distinct contributions for LP-derived signals in decision formation: (i) a pooled |ΔDir|-dependent signal that gates entry into an update regime ("Update Gate"), (ii) a |ΔDir|-dependent modulation of sensory gain that scales the influence of current evidence during updating ("Gain"), and (iii) an interhemispheric imbalance that biases choice direction upon updating, which can arise from either signed or unsigned LP comparison signals ("Bias/Reference"). Refer to the section on mechanistic specificity of optogenetic modulation for analyses testing which of these components must be directly perturbed to reproduce the optogenetic phenotype.

**Update gate as a latent variable:**
In the framework used here, p(Update) reflects entry into a latent internal state that changes how current sensory evidence is weighted, whereas p(Repeat) is a behavioral statistic defined over pairs of trials and depends on both internal state and the sequential structure of sensory evidence. Critically, entering the update state does not necessitate a switch in choice. Instead, updating reweights the decision variable, and when combined with a directional bias – such as that introduced by unilateral optogenetic perturbation – the resulting reduction in choice variability can manifest as an increase in p(Repeat) in spite of being in the p(Update) state more. When |ΔDir| is large and updating is frequent, unilateral perturbations exert their strongest influence, making choices more consistent across adjacent trials. This produces an increase in p(Repeat) that scales with |ΔDir|, even though the underlying probability of updating is also increasing. Thus, changes in p(Repeat) under optogenetic perturbation primarily reflect biased readout during update trials rather than changes in the likelihood of updating itself, and p(Repeat) should not be interpreted as a direct measure of update probability.

**Model validation:**
We first validated whether our reduced model operated in a regime consistent with normative task behavior. Because the model architecture explicitly introduces |ΔDir|-dependent updating, some dependence of performance metrics on |ΔDir| is expected by design. We therefore treat these baseline simulations as a validation step rather than as an independent prediction. Simulated choices reproduced |ΔDir|-dependent changes in psychometric slopes and the psychometric dependence on signed coherence (fig. S16A and B). We also observed the characteristic decrease in p(Repeat) with increasing |ΔDir| (fig. S16C), mirroring the empirical behavioral data (Fig. 1, figs. S2). Importantly, the model does not explicitly encode a tendency to repeat choices. The dependence of p(Repeat) on |ΔDir| emerges as a consequence of the model's trial-by-trial choice generation under update gating, and is treated here as a validation metric rather than a mechanistic control variable in the model. The model also recapitulated coherence-dependent trends in hit rate and repetition probability across |ΔDir| bins (fig. S2E,F), providing confidence that subsequent perturbation effects were grounded in a model capable of generating empirically observed behavioral patterns

**Comparison Signal Variants:** While the model design specifies how LP-derived signals influence updating and choice, it does not uniquely specify the form of the comparison signal itself. To assess which features of LP-derived comparison signals are constrained by the data, we examined several model variants that differed in how trial-to-trial sensory differences were represented prior to ACC readout.

Critically, across all variants, |ΔDir| signals are pooled and treated as unsigned for update gating and gain modulation. As a result, baseline performance and psychometric structure are largely preserved across variants (fig S16A-D), and differences emerge primarily in how optogenetic perturbations interact with the bias term.

Optogenetic manipulations that differentially affect gain, noise, or interhemispheric imbalance interact selectively with how directional information enters the decision variable, such that some combinations of comparison signal and perturbation mode produce lateralized effects while others do not. The simulations below accordingly focus on optogenetic conditions, where these interactions provide the strongest constraints on the form of LP-derived comparison signals

1. "Signed difference": LP preserves both the magnitude and sign of trial-to-trial ΔDirection, but remains anchored to sensory evidence rather than choice. This allows interhemispheric imbalance to bias choice direction upon updating without assuming choice encoding in LP.

2. "Unsigned comparator": LP encodes only the magnitude of sensory change |ΔDir|. In this case, the bias term lacks intrinsic directional information, such that lateralized effects can only arise when asymmetries are introduced through perturbation or downstream readout.

3. "Previous choice remapping": LP provides a signed ΔDir that is mapped to the animal's previous choice. This variant assumes choice-referenced information that is not supported by imaging data and therefore requires such a transformation to occur downstream, for example within ACC.

**Optogenetic Perturbation Modes**:
We next examined how different optogenetic perturbation modes interact with the model architecture. Because lateralized effects arise exclusively through the bias term, optogenetic perturbations provide a probe of how comparison signals are incorporated into the decision process, while still permitting substantial degeneracy across model variants.

All perturbations are implemented prior to interhemispheric subtraction and therefore manifest as changes in interhemispheric balance at the level of the bias/reference term that enters the decision variable.

1. “Gain”: Perturbations scale the amplitude of LP-derived comparison signals. This alters the magnitude of interhemispheric imbalance without changing the structure of the comparison signal itself, effectively amplifying or attenuating directional biases upon updating.

2. “Offset”: Perturbations introduce an additive offset to one hemisphere, directly imposing an interhemispheric asymmetry. This offset is combined with the sensory comparison signal at the level of the bias term, providing a structured asymmetry that biases choice direction regardless of the sign of the underlying comparison signal.

3. “Noise”: Optogenetic stimulation is modeled as degrading the endogenous |ΔDir| signals provided by LP, by replacing the signal with random noise. This tests the consequences of eliminating informative sensory comparison signals while preserving the overall model architecture.

**Constraints for model selection:**

To clarify how different forms of LP-derived comparison signals and optogenetic perturbations shape behavior within the model, we systematically evaluated combinations of comparison signal content and optogenetic transformation using a common architecture. **Table S5** summarizes whether each combination reproduces key qualitative features of the behavioral data under unilateral and bilateral perturbation, providing a checklist of constraints imposed by the experiments rather than a ranking of model performance.

**Constraint 1: Unilateral optogenetic stimulation must generate a directional choice bias.** Variants lacking an explicit interhemispheric imbalance term (A1), or in which perturbations entered primarily as noise (A3), failed to produce consistent lateralized effects.

**Constraint 2: Lateralized bias must scale with |ΔDir|.**
Several variants produced directional biases that were either insensitive to |ΔDir| or strongest at low |ΔDir| (A2 and B2), failing to capture the monotonic scaling observed experimentally. Thus, |ΔDir|-dependent bias places a stronger constraint than lateralization alone.

**Constraint 3: Bilateral perturbation must cancel directional bias while preserving performance.** Variants in which directional bias arose from interhemispheric imbalance naturally satisfied this condition, as symmetric perturbations increased comparator magnitude without introducing net directional bias. Models, however, varied in whether performance is preserved in bilateral stimulation. “Noise” variants (A3 and C3) globally impaired performance, but when regarded as unsigned comparator signals (B3), the impact was more limited to low

|ΔDir| where the noise increased the probability of updating when it is in fact more optimal to stay. For the same reason, if gain perturbation was applied at much higher values (not shown), we would see similar vulnerability of low |ΔDir| trials under bilateral stimulation.

Both unsigned comparison and previous-choice remapping variants were capable of reproducing the full set of behavioral signatures, but only under specific constraints on how interhemispheric imbalance modulated ACC activity.

Given that ACC promotes contralateral actions, the effective sign of imbalance in the decision variable determines whether perturbation produces excitation or suppression of contralateral action representations. Thus, the modeling constrains not the precise form of LP comparison signals, but the requirement that downstream ACC circuitry implements a sign-consistent transformation converting imbalance into directional bias.

**Mechanistic specificity of optogenetic modulation:**
In the models described above, optogenetic stimulation modifies the |ΔDir| term early upstream and thereby influences all 3 mechanisms (Update, Gain and Bias). To determine if we can recapitulate the same lateralized optogenetic effects by generating the bias term alone, we evaluated 3 variants of the architecture described above – focusing on the unsigned comparator variant with gain-based opto perturbation (B1), which provided the best qualitative recapitulation of our optogenetic results

First, in B1 (full mechanistic model, fig. S17A), optogenetic modification can influence all three computations (Update, Gain and Bias). In Model D ("OptoBias", fig. S17B), we retained the baseline architecture of computations that |ΔDir| contributes to. In this variant, update probability and gain modulation remained determined solely by the true |ΔDir|, and optogenetic stimulation affected only directional bias during updating. In Model E ("NoUpdate", fig. S17C), the mixture of stay-update regimes was removed entirely such that |ΔDir| entered the decision variable only through the bias term. This reduced architecture tests whether lateralized |ΔDir|-dependent bias can be reproduced without update gating or gain modulation.

Unilateral |ΔDir|-dependent lateralized choice biases were reproduced in both Models D and E (fig. S17B and C), indicating that modulation of the imbalance term is sufficient to account for the primary lateralized optogenetic phenotype. However, only models retaining update regime structure (Models B1 and D) reproduced the baseline non-monotonic psychometric sensitivity observed in normative behavior (fig. S16G, S17B).

We next generated simulated choices from each variant and fitted them using the same binomial GLMM applied to empirical data. In empirical data, two optogenetic interactions significantly improved model fit: DirCoh:Opto and |ΔDir|:OptoSide (Fig. 2P). In simulated datasets, inclusion of |ΔDir|:OptoSide improved fit across all variants, consistent with explicit implementation of |ΔDir|-dependent bias. Thus, in Models D and E, apparent slope differences arise primarily from |Δ|-dependent shifts rather than a global change in evidence sensitivity. Only the full mechanistic model (B1) recapitulated the contributions of both regression interactions simultaneously. Together, these analyses indicate that modulation of the imbalance term is sufficient to generate unilateral |ΔDir|-dependent lateralized biases, but reproducing the full set of empirical regression signatures requires perturbation of upstream comparison signals within the complete regime-based architecture.

**Other model predictions:**

Although bilateral LP-ACC stimulation did not produce lateralized choice biases or impair performance (fig. S8), the model makes specific predictions about altered internal computations. Because bilateral stimulation is symmetric, it does not generate a lateralized reference signal (fig. S16L), consistent with the data. However, it increases the pooled comparison magnitude that governs entry into the update regime, thereby increasing p(Update), particularly at low |ΔDir| (fig. S16J). This shift manifests as a non-monotonic change in p(Repeat) (fig. S16K): excessive entry into the update regime can impair performance at low |ΔDir| (eg. fig. S16U-X) , where staying is optimal, while potentially improving performance at intermediate or higher |ΔDir| by promoting appropriate switching. The magnitude of this effect depends on the specific mechanism (noise, offset, or gain), with gain-based perturbations producing the least disruption at low |ΔDir| (fig. S16I-K).

Nevertheless, although the model predicts increased p(Update) under bilateral stimulation, the behavioral consequences depend critically on each animal's intrinsic threshold for entering the update regime. In the model, update probability is governed by a nonlinear gate; bilateral stimulation shifts pooled comparison magnitude relative to this threshold. Animals operating below threshold may show enhanced appropriate switching at intermediate |ΔDir|, whereas animals already near saturation may exhibit impaired performance at low |ΔDir| due to excessive updating. Such heterogeneity can lead to weak or non-monotonic population-level effects, consistent with the modest trends observed when stratifying empirical data by |ΔDir| (fig. S4H). Thus, variability in update thresholds provides a mechanistic explanation for why bilateral perturbation does not produce a uniform behavioral signature.

**Summary:**

In this framework, LP-derived comparison signals encode trial-to-trial sensory change and regulate the probability of entering a latent update regime, but do not themselves encode action preference or repetition. Entry into an update regime reflects a computational shift from maintaining the previous choice toward re-assessing incoming evidence, and does not necessarily involve choice switching. Instead, it determines how |ΔDir| shapes the decision variable. Behavior reflects a mixture of "stay" and "update" regimes: when |ΔDir| is small, decisions are dominated by persistence of the previous choice, whereas larger deviations increase the likelihood of entering an update regime in which current sensory evidence is re-evaluated.

Directional biases emerge only when these comparison signals are read out through an interhemispheric opponent mechanism that converts imbalance into action preference. Our simulations show that the |ΔDir|-dependent lateralized biases we observed with unilateral optogenetic perturbations primarily acts through this bias term. Together, the modeling supports a circuit-level account in which LP conveys information about deviations from recent sensory evidence, while downstream circuits such as ACC implement the regime transitions and transformations necessary to translate those deviations into changes in behavior. This architecture provides a mechanism by which sensory history flexibly governs when to persist versus when to revise a choice.

**Supplementary Note 2: Formulation of the |ΔDir|-driven updating model**

The model was designed as a reduced, mechanistic framework linking LP-derived trial-to-trial sensory comparison signals to choice updating. Simulations operate on the empirical trial sequence and generate choices via Monte Carlo sampling.

On each trial t, the model computes:

1. A bilateral sensory comparison signal derived from the difference between consecutive stimuli
2. A comparison magnitude that governs the probability of entering an update regime
3. A regime-dependent decision variable used to generate choice

Choice probability is obtained by passing the decision variable through a logistic function, and binary choices are sampled from this probability.

**Stimulus representation:**
$x_t$ denote signed stimulus strength on trial t ranging from -1 to 1 with 8 unique values as in the task.

Signed or unsigned ΔDir information enters identically in parallel to each hemisphere as $\Delta d_L$ and $\Delta d_R$ indicating left and right hemisphere terms respectively. The sign of the raw signal depends on the variant.

In unsigned comparator, information enters as unsigned magnitudes

$|\Delta d_L|$ and $|\Delta d_R|$, each ranging from 0 to 2

In signed difference variant,

$\Delta d_L$ and $\Delta d_R$, each ranging from -2 to 2

In previous choice remapping,

$\Delta d_L$ x PrevChoice

$\Delta d_R$ x PrevChoice

where PrevChoice is +1 if right, -1 if left, anchoring the comparison signals to the previous choice before computing imbalance.

**Update gating**

The magnitude of comparison signals governs the probability of entering an update regime.

First, a pooled comparison magnitude is computed based on |ΔDir| signals from both hemispheres:

$$U_t = |\Delta d_{L(t)}| + |\Delta d_{R(t)}|$$

The probability of updating (switching regime) is given by a logistic function:

$$p(\text{Update})_t = \text{sigmoid}(\ \text{beta}^{\text{update}} \times (\ U_t - T^{\text{update}}\ )\ )$$

where:

- $\text{beta}^{\text{update}}$ controls the steepness of the transition, and was set as 8 in all simulations.
- $T^{\text{update}}$ is the update threshold, set as 0.6 for all simulations.

On each trial, the model enters the update regime with probability $p(\text{Update})_t$ and otherwise remains in the stay regime.

**Bias/Reference Signal**

The model constructs bilateral comparison channels $\Delta d_{L(t)}$ and $\Delta d_{R(t)}$. The reference term always represents an interhemispheric imbalance in LP-derived comparison signals.

In unsigned comparator:

$$\text{ref}_t = |\Delta d_{R(t)}| - |\Delta d_{L(t)}|$$

In signed difference and previous choice remapping:

$$\text{ref}_t = \Delta d_{R(t)} - \Delta d_{L(t)}$$

Thus, the reference signal always captures the relative strength of right versus left comparison signals.

**Decision Variable**

The model uses two decision variables (DV), one for the stay regime and one for the update regime.

Stay regime DV:

$$DV_{\text{stay},t} = k * x_t + \text{ref}_t$$

Update regime DV:

$$DV_{update,t} = (k/4 + \Delta z_t^{gain}) * x_t + ref_t$$

Where k is the sensitivity to evidence (x), and where the pooled comparison magnitude:

$$\Delta z_t^{gain} = (|\Delta d_{R(t)}| + |\Delta d_{L(t)}|)/4$$

Both DVs incorporate the bias term ($ref_t$).

The key difference between regimes lies in the gain on sensory evidence $x_t$. In the stay regime, gain remains high (k), consistent with strong performance when consecutive stimuli are similar. In the update regime, gain is reduced to k/4, reflecting a cost associated with switching away from the previous decision state. However, this reduced baseline gain is partially offset by $\Delta z_t^{gain}$, which increases with the magnitude of comparison signals. Thus, stronger trial-to-trial sensory differences increase the effective weighting of sensory evidence within the update regime.

**Choice probability:**

Choice probability is computed via logistic readout of the DV:

$$P(ChooseR)_t = sigmoid(DV_t)$$

A binary choice is then sampled:

$$Choice_t \sim Bernoulli(P(ChooseR)_t)$$

**Optogenetic perturbations:**

Perturbations were implemented as transformations of $\Delta d_{L(t)}$ and/or $\Delta d_{R(t)}$. prior to computing $U_t$ (update), $ref_t$ (bias) and $\Delta z_t^{gain}$ (gain).

For gain:

$$\Delta d_t = g * \Delta d_t$$

where g scales the comparison signal (g = 2 in simulations)

For offset:

$$\Delta d_t = \Delta d_t + r$$

where r is an additive offset (r = 0.5 in simulations)

For noise:

$$\Delta d_t = \varepsilon_t \sim \text{Uniform}(-q, q)$$

Where q sets the range of noise values, set to be 1 in all noise simulations.

**Supplementary Figure 1**

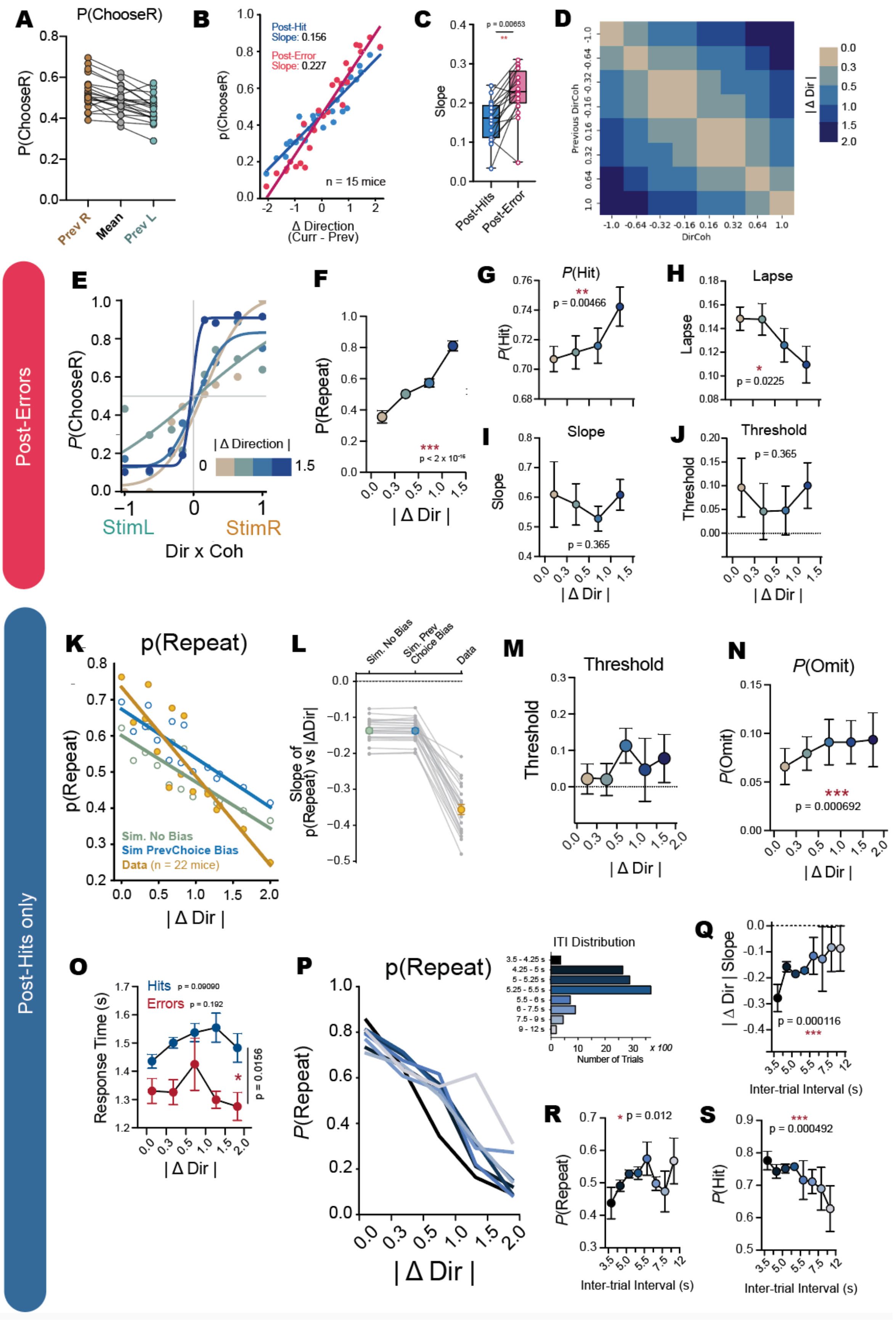

**Supplementary Figure 1: |ΔDir|-dependent behavioral performance during 2AFC RDK direction discrimination task**

(**A**) Probability of choosing right (P(ChooseR) based on previously rewarded choice, matched within mice (n = 22), showing tendency to repeat previously rewarded choice. (**B**) P(ChooseR) as a function of signed ΔDirection (current − previous stimulus direction), shown separately for post-hit and post-error trials. Linear fits indicate significant dependence on ΔDirection, with steeper slopes following errors. (**C**) Comparison of ΔDirection slopes from (A), showing enhanced sensitivity to trial-to-trial sensory change following errors. Each dot represents 1 mouse. N = 15 mice. (**D**) Heatmap with colors representing the |ΔDir| bin that each combination of current and previous stimulus direction fell into. (**E**) Psychometric functions for post-error trials, stratified by |ΔDir|. (**F** to **J**) (F) P(Repeat), (G) Hit rate, (H) Lapse rate, (I) Psychometric slope and (J) Psychometric threshold, as a function of |ΔDir|. Only bins up to 1-1.5 |ΔDir| shown as there were limited trials in the final 1.5-2.0 |ΔDir| bin that were after errors.  showing a modest decrease with increasing trial-to-trial sensory change. (**K** and **L**) Simulation results, comparing data to a model sampling only from mean P(ChooseR) (Sim. No Bias) and a model where P(ChooseR) depended on previous choice (Sim PrevChoice Bias). The dependence of P(Repeat) on |Δ Dir| in the data (yellow) is steeper than expected from behavioral models with no bias (green) or simple previous choice biases that does not regard strength of relative evidence (blue), suggesting that relationship between P(Repeat) and |ΔDir| is not a simply an artfact of consecutive stimulus regularities. (K) Comparison of slopes in (J) on a per mouse basis (n = 22 mice). Results reported only for previous hits. (**M**) Psychometric Threshold as a function of |ΔDir|. (**N**) P(Omit) as a function of |ΔDir|. (**O**) Influence of |ΔDir| on reaction times.  Reaction times were faster in error trials. The influence of | Δ Dir | on response times on hit and error trials followed an inverted-U shape trend, being fastest in trials of least ambiguity in |ΔDir| (the poles: most similar - 0, and most different - 2) and slower in the face of more ambiguous relative evidence, and was not significant in a linear model (hits:p = 0.09090, errors: 0.192). (**P**) The influence of |ΔDir| on P(Repeat) of rewarded trials, plotted for trials binned by duration of ITIs. (**Q**) The strength of |ΔDir| influence (negative slope) is greatest with short ITIs, and decays with longer ITIs. p-value reported for the effect of ITI on slope describing the relationship between | Δ Dir | and P(Repeat) (as in o), repeated measures ANOVA. (**R**) Effect of ITI on P(Repeat), p-value reported for variance explained by continuous values of ITIs in binomial GLME of P(Repeat) which accounts for previous and current trial difficulty. (**S**) P(Hit) was greater with shorter ITIs. p-value reported for variance explained by continuous values of ITIs in binomial GLME for P(Hit) which accounts for previous and current trial difficulty.

**Supplementary Figure 2**

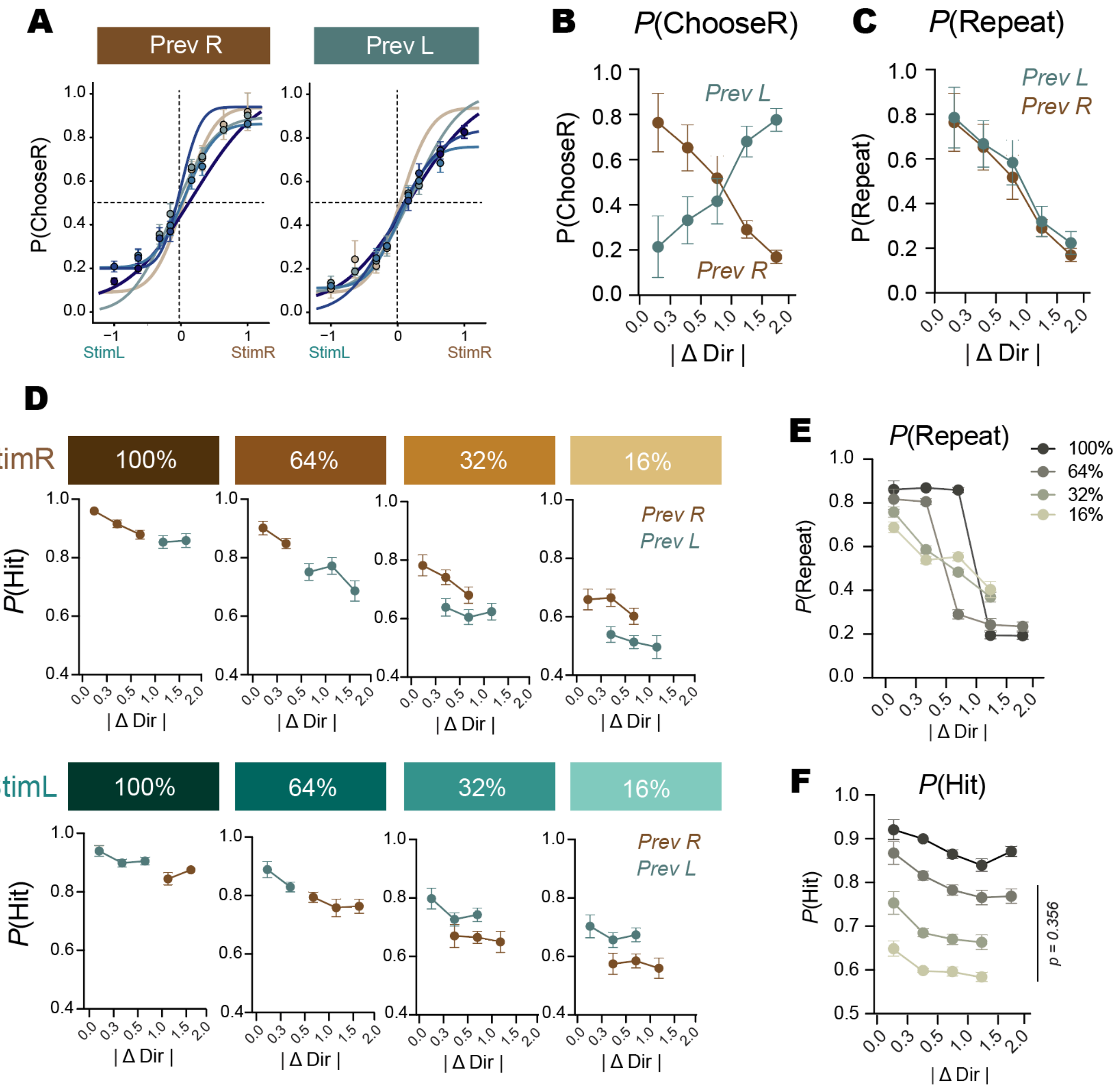


**Supplementary Figure 2: |ΔDir|-dependent behavioral effects are symmetric with respect to previous choice**

(**A**) Psychometric functions (P(ChooseR)) plotted separately for trials following a previous right (PrevR) or previous left (PrevL) choice. (**B**) P(ChooseR) as a function of |ΔDir|, separated by previous choice. Choice probability shifts systematically with |ΔDir| in different directions for PrevR and PrevL reflecting symmetric history effects. (**C**) Probability of repeating the previous choice, P(Repeat), as a function of |ΔDir| for PrevR and PrevL trials. Repeat probability decreases monotonically with |ΔDir| in a symmetric manner for both previous choice directions. (**D**) Hit rate, P(Hit), as a function of |ΔDir| for rightward (StimR) stimuli, shown separately by stimulus direction and coherence, and colored by previous rewarded choice. Performance improves with increasing |ΔDir| across coherence levels. (**E**) P(Repeat) as a function of |ΔDir| plotted separately for different stimulus coherences, illustrating that |ΔDir|-dependent history effects are present across stimulus strengths. (**F**) P(Hit) as a function of |ΔDir| separated by stimulus coherence. across stimulus direction and coherence, showing that |ΔDir| impacts performance accuracy similarly across different stimulus strengths.

**Supplementary Figure 3**

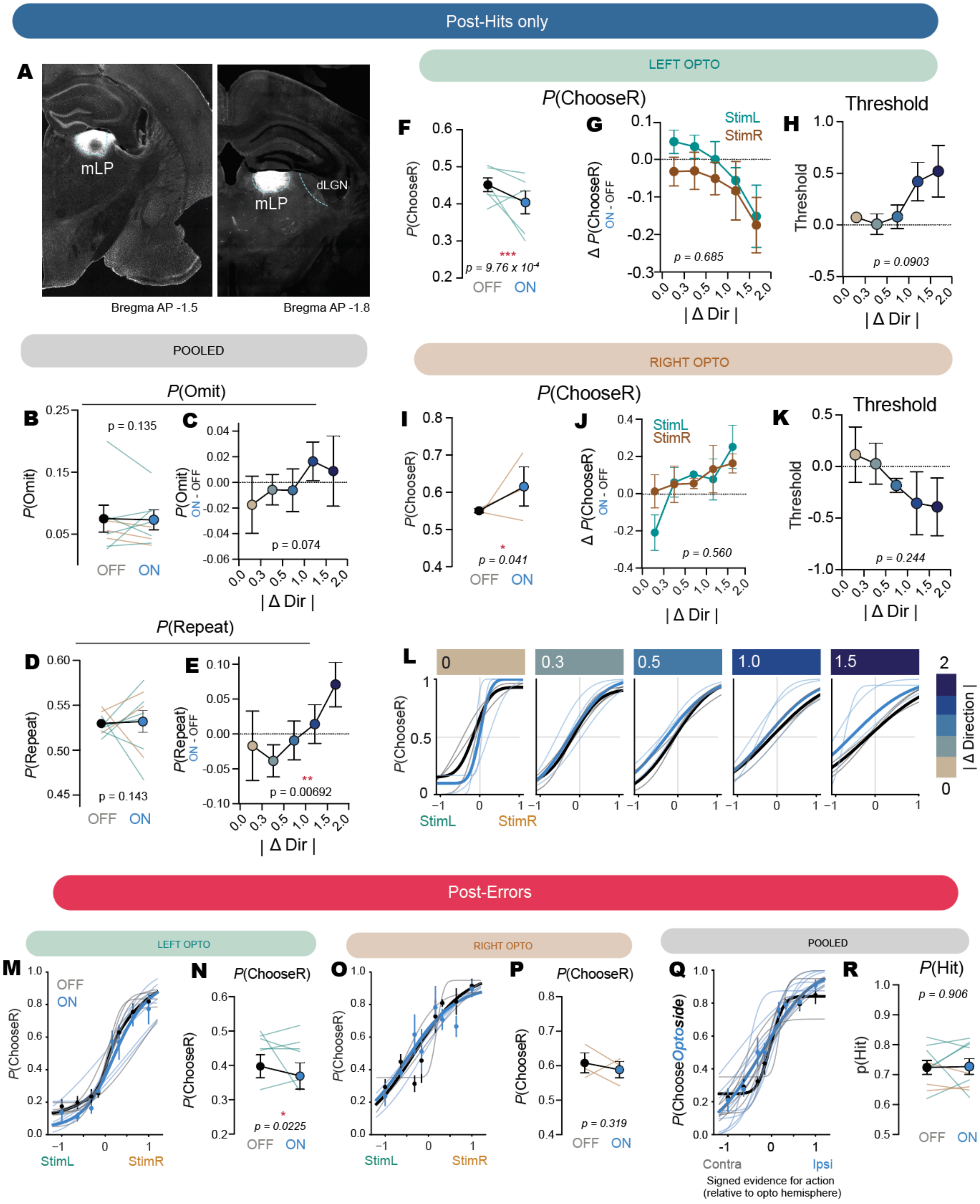

**Supplementary Figure 3: Effects of unilateral LP-ACC optogenetic stimulation across additional behavioral metrics**

(**A**) Representative histology from two different animals showing ChR2 expressing restricted to medial LP. (**B**) Probability of omitting a trial (P(Omit)) in light OFF and light ON trials, shown for individual animals, turquoise indicates left- and brown indicates right-stimulation cohort. Population average shown pooled across n = 9 animals. (**C**) P(Omit) as a function of |ΔDir|. (**D**) Probability of repeating a previously rewarded trial (P(Repeat)) in light OFF and light ON trials, shown for individual animals, turquoise indicates left- and brown indicates right-stimulation cohort. (**E**) P(Repeat) as a function of |ΔDir|. Note that this can be secondary consequence of increased choice bias leading to reduced choice variability and thus higher P(Repeat). (**F**) Probability of choosing right in light OFF and light ON trials for left LP-ACC axonal stimulation, shown for individual animals. (**G**) Change in P(ChooseR) (ON-OFF) as a function of |ΔDir|, showing no difference or interaction when stratified by stimulus direction. (**H**) Fitted psychometric thresholds (here equivalent to point of subjective equality) as a function of |ΔDir| trended towards increase with |ΔDir| but was not statistically significant. (**I** to **K**) Same as (F) to (H) but  for right LP-ACC stimulation. (**L**) Psychometric functions for right LP-ACC OFF and ON stimulation trials, stratified by |ΔDir| (absolute change in stimulus direction from the previous trial). (**M**) Psychometric functions (P(ChooseR)) for left LP-ACC opto-stimulation trials (ON; blue) and interleaved control trials (OFF; black). Only for previous errors. Thin lines indicate individual animals; thick lines indicate population mean. (N) Probability of choosing right in light OFF and light ON trials for left LP-ACC axonal stimulation, shown for individual animals, for previous errors. (**O** and **P**) Same as (M and N) but for right LP-ACC opto-stimulation trials. (Q) Psychometric functions aligned to optogenetic light stimulation hemisphere, pooled across left and right stimulation cohorts. Only for previous errors. (**R**) Probability of hit (P(Hit) in light OFF and light ON trials. Averaged across left- and right-stimulation cohorts. Color of thin Individual lines indicate stimulation cohort (turquoise for left and brown for right). All binary trial level statistics derived from binomial GLMMs. Thresholds estimated with linear mixed effects models.

**Supplementary Figure 4**

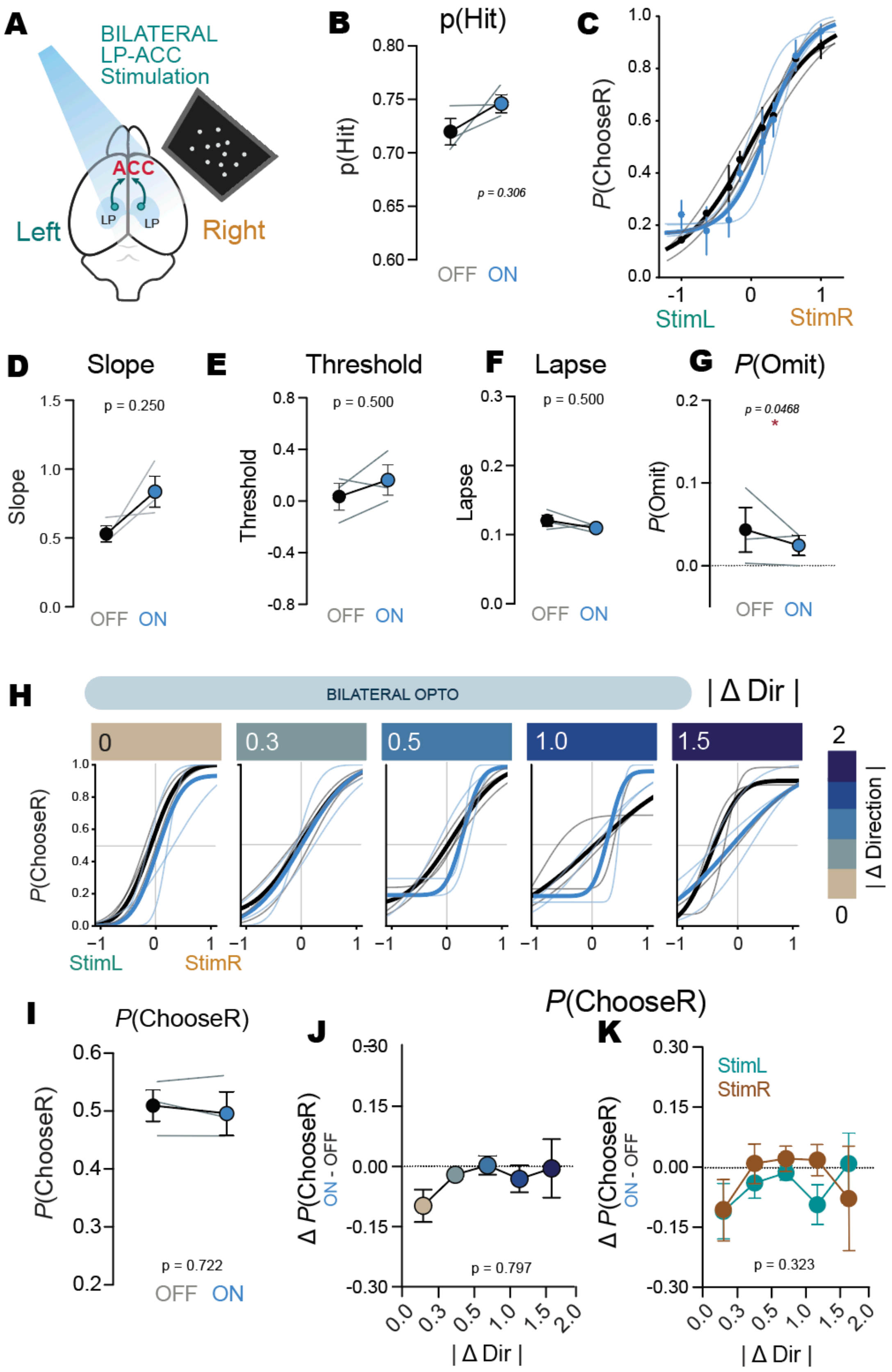

**Supplementary Figure 4: Bilateral LP-ACC stimulation cancels out lateralized biases and preserves performance**

(**A**) Experimental schematic for bilateral stimulation experiment. AAV carrying ChR2 was injected bilaterally in medial LP, and their axons illuminated over the same window along the midline in ACC. (**B**) Probability of a hit (P(Hit)) for light OFF and light ON trials, shown for n = 3 individual mice. (**C**) Psychometric fits with (light ON, blue) and without (light OFF, black) optogenetic activation in the previous trial. Bold population fits, thin lines individual animals. (**D**) Psychometric function slopes trended steeper across all animals but was not statistically significant. (**E**) No consistent shift in threshold across animals (F) Lapses remained unchanged. (**G**) Probability of omissions (P(Omit)) was reduced for light OFF and light ON trials, shown for n = 3 individual mice. D to F two-tailed Wilcoxon signed-rank test. (**H**) Psychometric functions for bilateral LP-ACC stimulation ON and OFF trials, stratified by |ΔDir|. (**I**) Probability of ChooseR (P(ChooseR)) for light OFF and light ON trials was unchanged by bilateral LP-ACC stimulation, indicating no lateralized biases. (**J** and **K**) Change in ΔP(ChooseR) (ON - OFF) stratified by |ΔDir| (J) or further separated into stimulus direction (K). Unlike unilateral stimulation, bilateral stimulation did not lead to |ΔDir|-dependent lateralized biases. B, G and I to K p-values from binomial GLMMs (Table S2).

**Supplementary Figure 5**

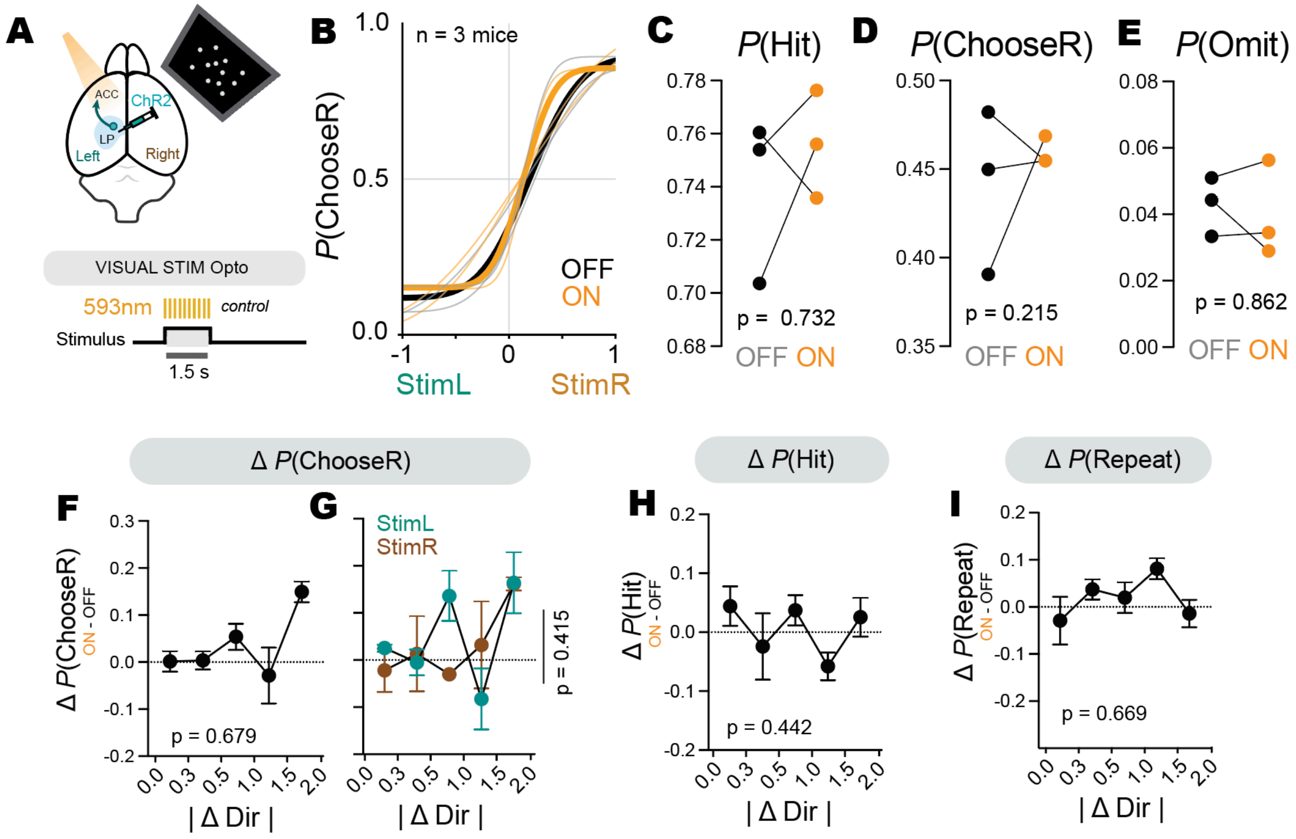


**Supplementary Figure 5: Illuminating ChR2-expressing LP-ACC axons with 593nm light had no effect on behavior performance.**

**(A)** Experimental schematic for control experiment with 593nm light delivered at the same frequency (20 Hz) as with blue light in the true experiment, during the stimulus presentation epoch. **(B)** Psychometric fits for performance during laser off and laser on trials for n = 3 mice. **(C to E)** Illuminating LP-ACC axons with 593nm light did not have a significant impact on (C) P(Hit), (D) P(ChooseR), or (E) P(Omit). Each averaged value for a single mouse samples equally from all stimulus levels. **(F to I)** 593nm light also did not influence |ΔDir|-dependent changes in (F) P(ChooseR) generally or (G) when stratified by stimulus direction. (H) No change in P(Hit) as a function of |ΔDir|. (I) No change in P(Repeat) as a function of |ΔDir|. C to I p-values from binomial GLMMs (Table S2).

**Supplementary Figure 6**

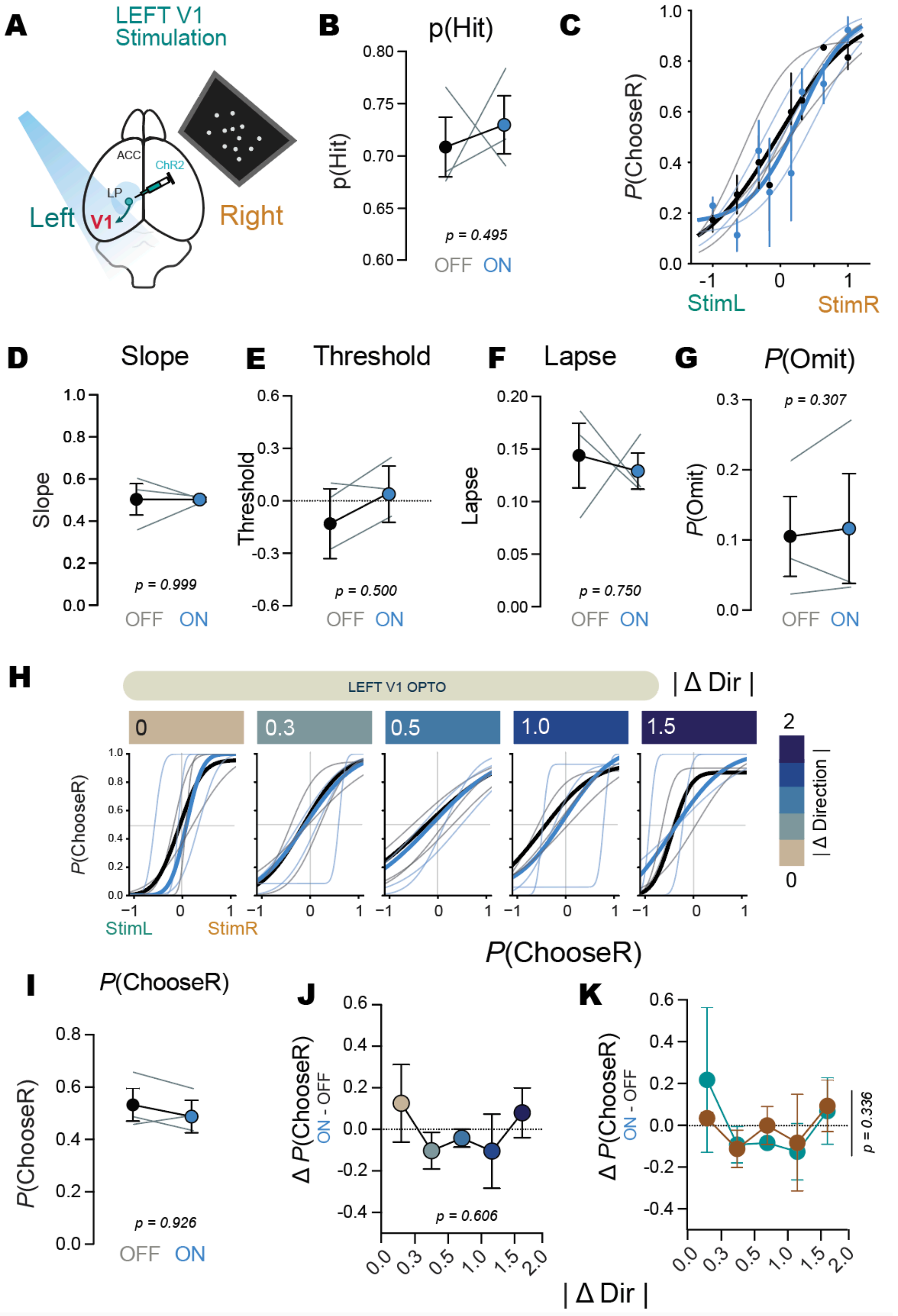

**Supplementary Figure 6: LP-V1 (left) stimulation did not lead to lateralized biases observed in LP-ACC stimulation**

(**A**) Experimental schematic for LP-V1 stimulation experiment. AAV carrying ChR2 was injected into left medial LP, in the same LP subregion targeted in all experiments. Their axons were activated by blue light illumination over V1.. (**B**) Probability of a hit (P(Hit)) for light OFF and light ON trials, shown for n = 3 individual mice. (**C**) Psychometric fits with (light ON, blue) and without (light OFF, black) optogenetic activation in the previous trial. Bold population fits, thin lines individual animals. (**D)** Psychometric slopes were not affected by LP-V1 stimulation, indicating preserved perceptual sensitivity (**E**) No consistent shift in threshold across animals. (**F**) Lapses remained unchanged. D to F two-tailed Wilcoxon signed-rank test. (**G**) Probability of omissions (P(Omit)) was unchanged for light OFF and light ON trials, shown for n = 3 individual mice. (**H**) Psychometric functions for bilateral LP-ACC stimulation ON and OFF trials, stratified by |ΔDir|. (**I**) Probability of ChooseR (P(ChooseR)) for light OFF and light ON trials was unchanged by bilateral LP-ACC stimulation, indicating no lateralized biases. (**J** and **K**) Change in ΔP(ChooseR) (ON - OFF) stratified by |ΔDir| (J) or further separated into stimulus direction (K). Taken together, unilateral stimulation of LP-V1 axons did not recapitulate lateralized or |ΔDir|-dependent behavioral effects of LP-ACC unilateral stimulation, in spite of the same LP region being targeting, suggesting target specificity of LP-ACC stimulation effects. B, G and I to K p-values from binomial GLMMs (Table S2).

**Supplementary Table 7:**

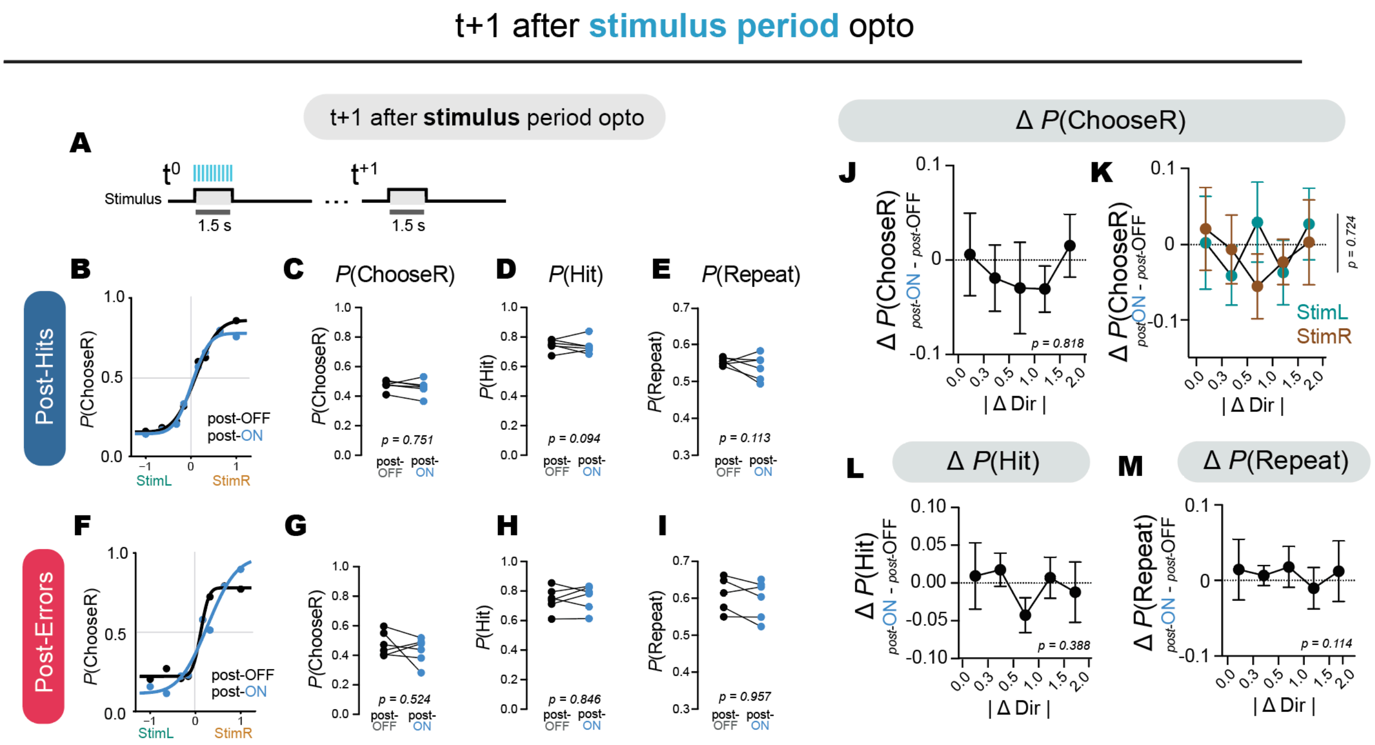


**Supplementary Figure 7: Optogenetic activation during the visual stimulus evaluation epoch has no impact on the subsequent trial.**
(**A**) Schematic showing optogenetic activation period during the visual stimulus period in the prior trial (t0). This is the same experiment as described in Fig 2, but considering the impact of optogenetic stimulation on the next trial. (**B**) Psychometric fits with (blue) and without (black) optogenetic activation in the previous trial. (**C** to **E**) No change in (C) P(ChooseR), (D) P(Hit), (E) P(Repeat) after hit trials with optogenetic stimulation. (**F** to **I**) Same as (C to E) but after error trials. (**J** to **M**) We found no disruption in the relationship between |ΔDir| and P(ChooseR) in across all stimuli (J) and stratified by stimulus direction (K). No significant interaction between previous opto-stimulation trial and |ΔDir| on (L) P(Hit), or (M) P(Repeat). p-values indicated in C to E, and G to M from binomial GLMMs (Table S2).

**Supplementary Figure 8:**

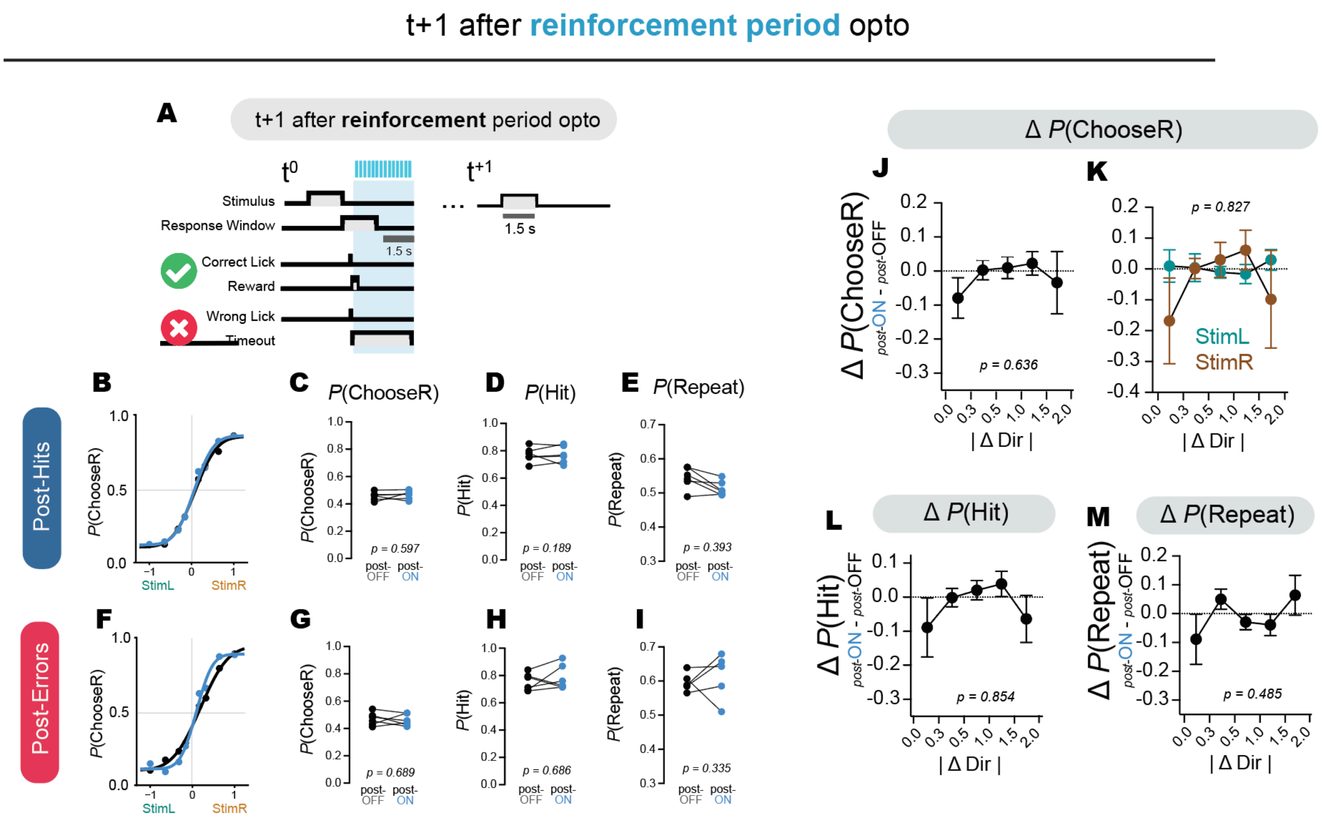


**Supplementary Figure 8: Optogenetic activation during the reinforcement epoch has no impact on the subsequent trial.**
(**A**) Schematic showing optogenetic activation period during the reinforcement period in the prior trial (t0). We examine the effect on the trial after optogenetic stimulation. (**B**) Psychometric fits with (blue) and without (black) optogenetic activation in the previous trial. (**C** to **E**) No change in (C) P(ChooseR), (D) P(Hit), (E) P(Repeat) after hit trials with optogenetic stimulation. (**F** to **I**) Same as (C to E) but after error trials. (**J** to **M**) We found no disruption in the relationship between |ΔDir| and P(ChooseR) in across all stimuli (J) and stratified by stimulus direction (K). No significant interaction between previous opto-stimulation trial and |ΔDir|on (L) P(Hit), or (M) P(Repeat). p-values indicated in C to E, and G to M from binomial GLMMs (Table S2).

**Supplementary Figure 9**

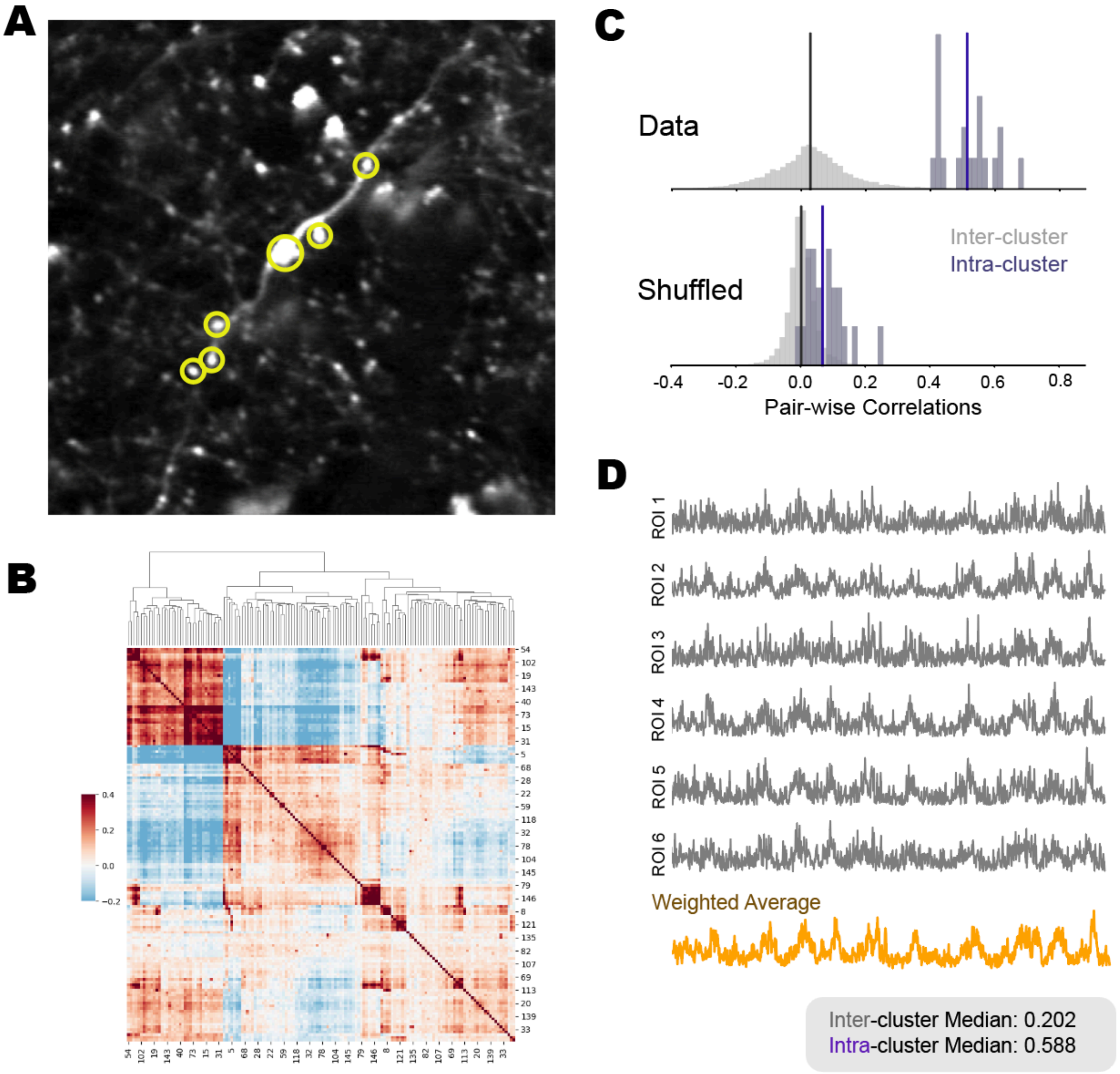


**Supplementary Figure 9: Processing steps in clustering individual bouton ROIs into axonal units.** (**A**) Maximum Projection of a representative field of view. Axonal boutons found to be correlated as a cluster are highlighted. (**B**) Correlation matrix for each axonal bouton in the field of view, with every other bouton. This correlation matrix then undergoes hierarchical clustering based on their similarity. (**C**) An empirical threshold is determined for each session/field-of-view such that the resulting axon clusters have a threshold minimum for intra-cluster correlation, and maximal inter-cluster correlation. (**D**) Raw traces (before denoising) for each axonal bouton identified in a cluster (highlighted in (A)). A weighted (by their signal-to-noise ratio) average of these boutons determined to belong in a cluster is taken to represent a single axonal cluster.

**Supplementary Figure 10**

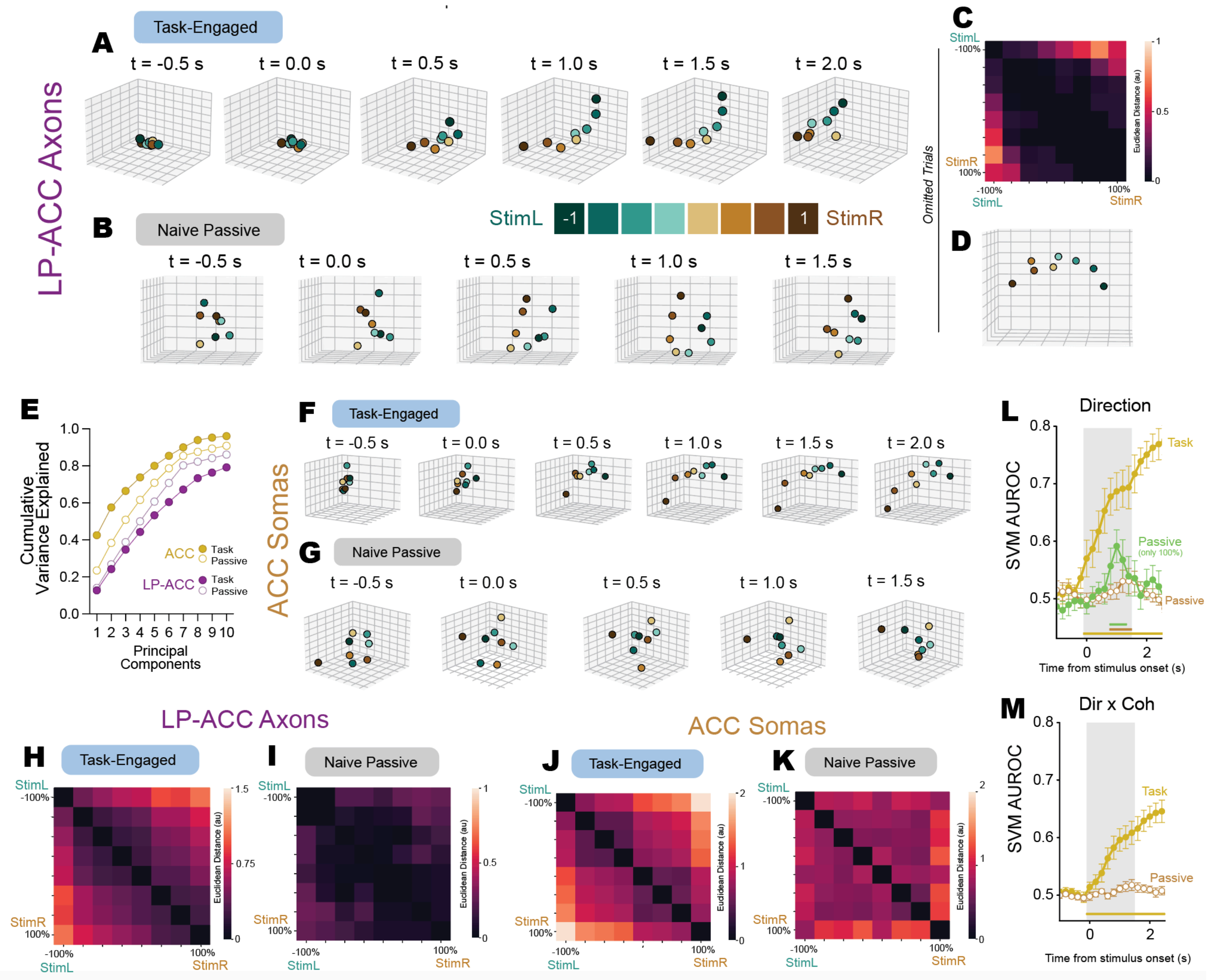


**Supplementary Figure 10: Task engagement reorganizes population representation of visual stimuli in a low-dimensional curved manifold.**
(**A** and **B**) LP-ACC populations averaged by stimulus signed coherence projected onto the first 3 principal components, during task engagement (A) and under naive passive viewing (B), visualized across different timepoints throughout stimulus presentation (1.5s in both cases). Population activity evolves across the stimulus presentation to form a low dimensional representation of each unique stimulus along a curved manifold. Along this curved manifold, every stimulus is represented with reference to the decision variable. Under passive viewing, the

population activity shows brief direction discriminability, with much less structure. Both low dimensional representations were derived from PCA on a trial- averaged activity across a pseudo-population, pooling all sessions. (**C**) Pair-wise Euclidean distances (arbitrary units) in the top 3 PCs between stimuli for omitted trials. (**D**) LP-ACC population projection for stimuli in omitted trials, at 1.5s after stimulus onset timepoint. (**E**) Cumulative variance explained for the first 10 principal components for both LP (purple) and ACC (yellow) populations under task-engaged (filled) and naive passive viewing (open) conditions. (**F** and **G**) Same as A and B but for ACC populations. All pair-wise euclidean distances were taken from a window. (**H** to **K**) Pair-wise Euclidean distances (arbitrary units) in the top 3 PCs between stimuli for LP-ACC axons (H and I) and ACC (J and K) under task engagement (H and J) and passive viewing (I and K). (**L** and **M**) SVM decoding accuracy for ACC neurons for direction (L) and Dir x Coh (signed coherence) (M) during task (yellow) and under passive viewing (open brown circles). For direction decoding (L), we also performed decoding with only 100% coherence stimuli under passive viewing (green). Horizontal lines at the bottom of the plot indicate timepoints where SVM AUROC is statistically greater than AUROC from a decoder trained on shuffled labels (one-tailed paired t-test within session for each timepoint, $p<0.05$). Task-engaged: $n = 9$ sessions from 3 mice. Passive: $n = 14$ sessions from 3 mice.

**Supplementary Figure 11**

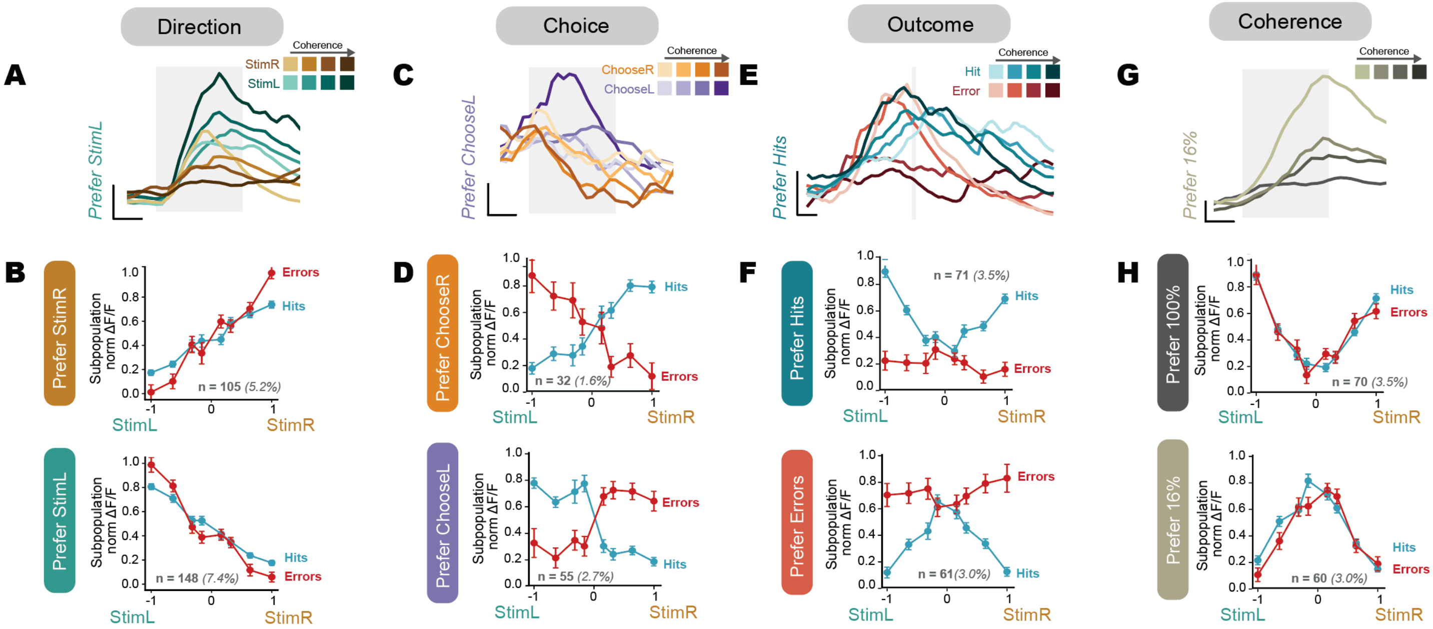


**Supplementary Figure 11: Subpopulation of LP-ACC axons with strong selectivity and monotonic scaling with coherence relative to task variables.**
Axons highly selective and with monotonic scaling for coherence graded task variables were identified with Kendall's Tau correlation using mean responses from 0.5s to 1.5s after stimulus onset. An axon was considered significantly modulated if the absolute Kendall's Tau obtained for a specific task variable was statistically greater ($p>0.05$, one-tailed t-test) than a null control set where signed coherence for the task variable was shuffled. To isolate proportions of axons that exhibit pure selectivity with monotonic scaling, we ensured that these subpopulations are non-overlapping, using their relative responses to hit and error trials helping to dissociate between highly correlated task variables such as direction and choice selectivity, as well as outcome and coherence. (**A** and **B**) Direction selectivity. (A) Trial-averaged traces of a representative StimL-prefering axon. (B) Direction-selective axons that preferred stimulus associated with right (top) and left (bottom) choice as a function of signed stimulus evidence. Errors and hits had similar responses as these axons showed greater fidelity to the true stimulus. (**C** and **D**) Choice selectivity. (C) Trial-averaged traces of a representative ChooseL-prefering axon (D) Choice-selective axons that preferred right (top) and left (bottom) side evidence as a function of signed stimulus evidence. Errors and hits had a crossed profile when plotted at a function of signed stimulus evidence, as axons reflected the absolute strength of evidence when selecting a side, whether or not evidence supported the choice. (**E** and **F**) Outcome selectivity. (E) Trial-averaged traces of a representative hit-prefering axon, grey line marks the time of reinforcement, which occurs 100ms after choice. (F) Outcome-predictive axons that preferred hits (or high evidence for choice taken) (top), and errors (or low evidence for choice) (bottom) as a function of signed

stimulus evidence. Errors and hits had mirrored v-shaped profiles when plotted at a function of signed stimulus evidence. The same stimulus was associated with different responses based on the outcome but in a way that is not dependent on the choice side, as the responses were vertically symmetric regardless of the stimulus or choice. (**G** and **H**) Coherence selectivity. (G) Trial-averaged traces of a representative low coherence-prefering axon. (H) Axons that preferred high coherence (100%, top), and low coherence (16%, bottom) as a function of signed stimulus evidence. In spite of showing a similar vertically symmetric v-shaped profile as f, these axons did not discriminate between outcomes. For A, C, E and G, all vertical scale bars correspond to 0.1 z-scored dFF, horizontal scale bars correspond to 0.5s. For A, C and G, the shaded area indicating the stimulus period.

**Supplementary Figure 12**

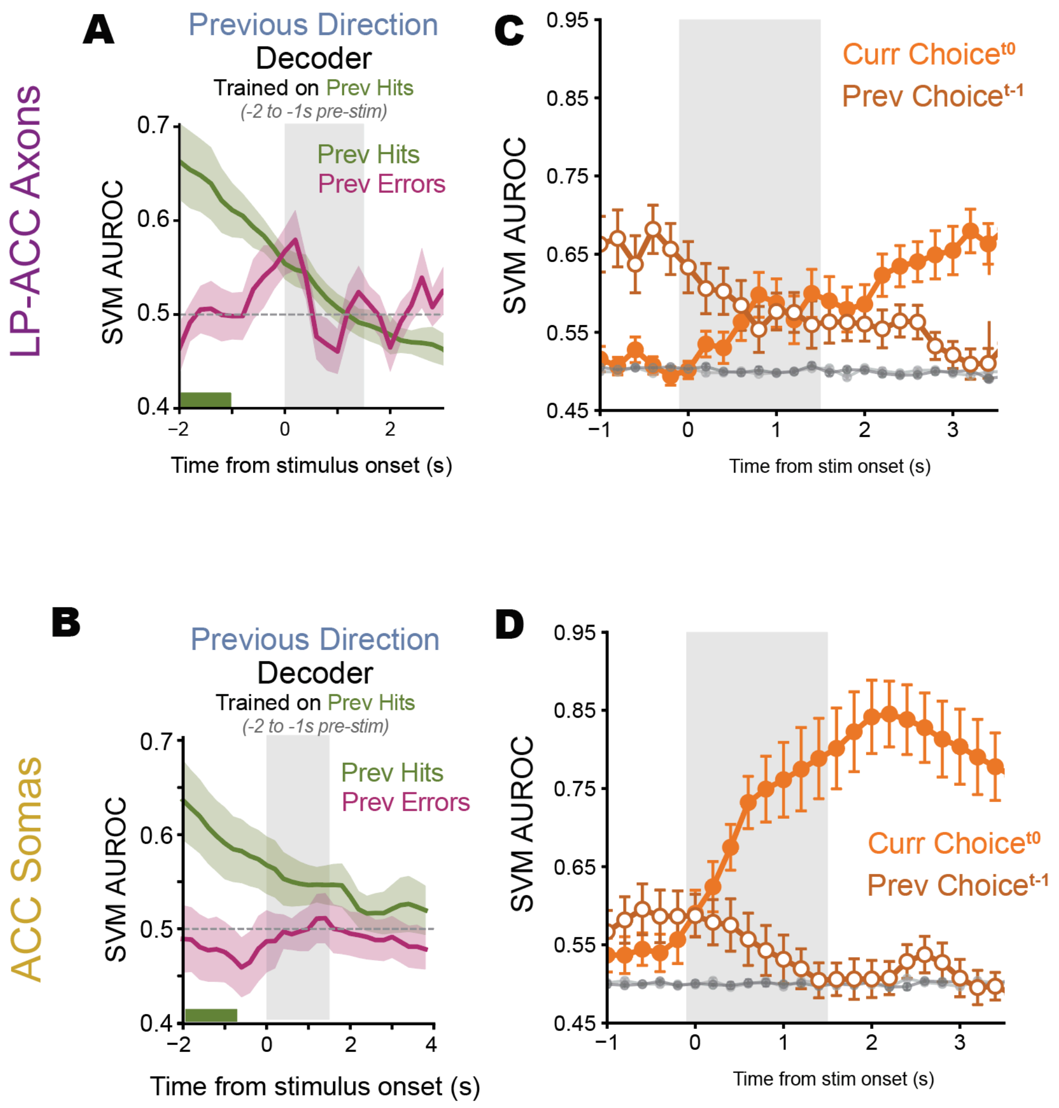


**Supplementary Figure 12: Cross-outcome decoding reveals stimulus-dominant history coding in LP–ACC but choice-dominant coding in ACC**

(**A**) Cross-outcome SVM decoding of previous direction with a training set composed of only previous hits. Decoder was trained on an averaged window -2 to -1s before stimulus onset (indicated by bottom bold green line), and then tested on held-out previous hit and previous error trials across all timepoints (100ms bins). As previous direction is identical to previous choice in all previous hit trials, using the same decoder to predict previous direction for previous error trials would succeed if LP-ACC axons predominantly carried previous direction (stimulus history) but result in SVM AUROC below 0.5 (wrong prediction) if LP-ACC axons predominantly carried previous choice. Notably, the cross-outcome decoder was able to generalize and decode previous direction above chance comparably for previous hits and errors during the 1s before stimulus onset (this pre-stimulus window excludes motor confounds because mice were required to withhold licking to initiate the next trial), and up to 500ms after stimulus onset. (**B**) Same as (A) but for ACC neurons. However, here the decoder failed to generalize to previous error trials for the prediction of previous direction, suggesting that previous-direction representations in ACC do not generalize across outcomes. (**C**) Linear SVM AUROC from decoding of current (filled) and previous (open) choice in LP-ACC axonal populations across time. Previous choice decoding declines from its peak during the ITI as a new trial begins. In LP-ACC axons, current and previous choice achieved comparable decoding performance during the stimulus evaluation period. (**D**) Same as (C) but for ACC neurons. Previous choice decoding similarly declined from its peak during the ITI. However, when a new trial begins, current choice information rapidly dominates ACC population representations. LP-ACC: n = 17 sessions from 5 mice. ACC: n = 9 sessions from 3 mice.

**Supplementary Figure 13**

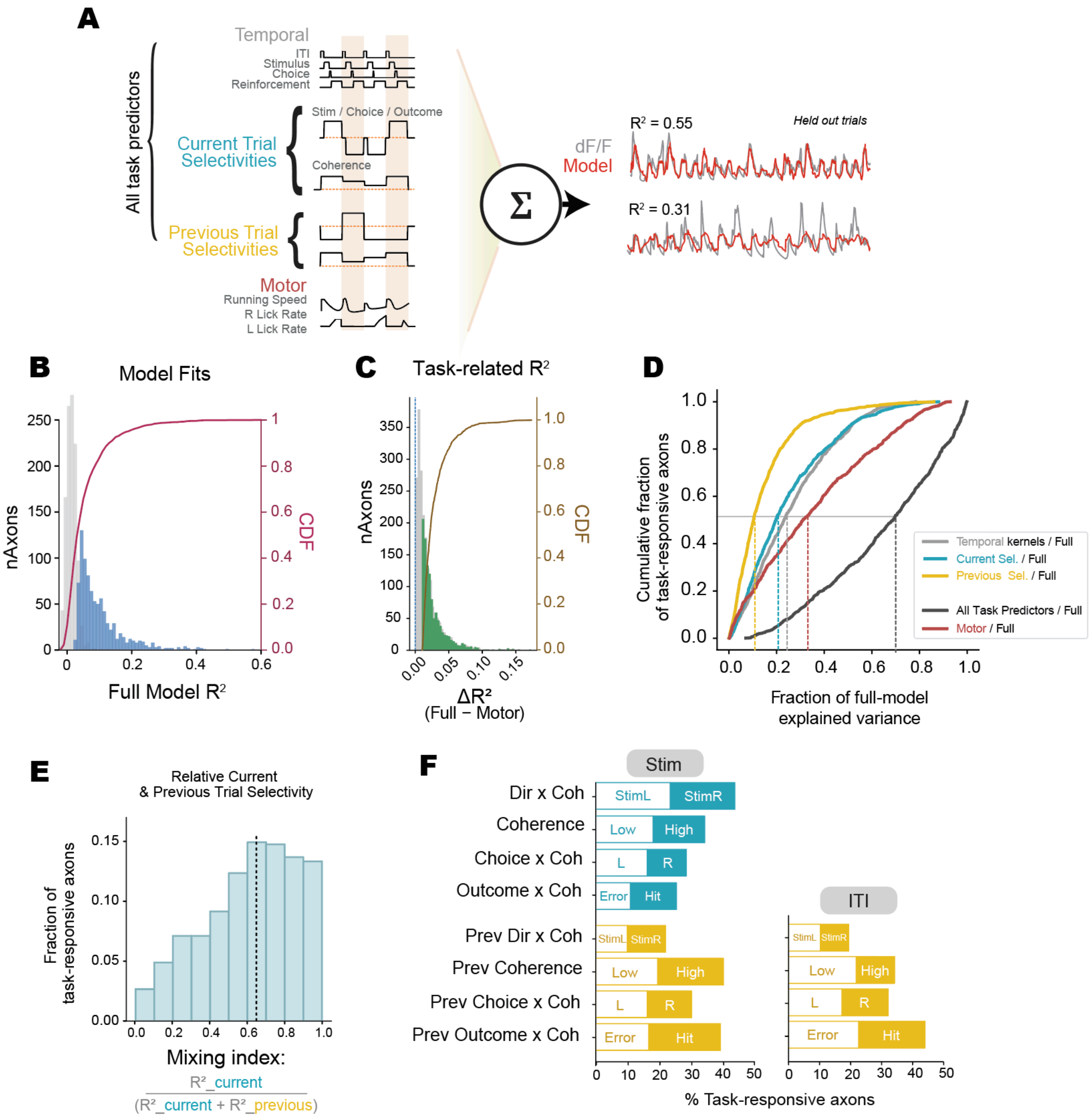

**Supplementary Figure 13: Single LP-ACC axon linear encoding models reveals robust task selectivity after accounting for motor-related variance**

(**A**) Schematic of the linear encoding model. The design matrix included temporal kernels, current-trial task variables (stimulus direction, coherence, choice, outcome), previous-trial variables, and motor-related predictors (running and licking; see Materials and Methods). Single-axon activity ($\Delta F/F$) was reconstructed as a linear weighted sum of all predictors. Example axons show observed activity (grey) and model reconstruction (orange). (**B**) Distribution of full-model cross-validated $R^2$ values across axons. Blue bars indicate task-responsive axons with reliable model fits on held-out data across 100 iterations (see Methods). (**C**) Distribution of task $R^2$, computed as full-model $R^2$ minus variance uniquely explained by motor predictors. Substantial task-related variance remains after accounting for motor-related variance. (**D**) Cumulative distribution of the fraction of full-model explained variance attributable to each predictor group across task-responsive axons. For each axon, the contribution of temporal kernels, current-trial variables, previous-trial variables, motor predictors, and all task predictors combined was computed relative to the full-model $R^2$. Dashed lines indicate medians. Although motor-related predictors account for the largest share of explained variance across the population, task predictors, particularly current-trial variables, explain substantial and non-trivial fractions of variance in a large proportion of axons. (**E**) Distribution of mix index among task-responsive axons. The mix index quantifies the relative contribution of current- versus previous-trial task predictors (excluding temporal kernels), where 0 indicates exclusively previous-trial selectivity and 1 indicates exclusively current-trial selectivity. Median mix index = 0.62, indicating dominant current-trial encoding with substantial multiplexing of past information. (**F**) Proportion of task-responsive axons with significant modulation by each predictor, and the specific task variable they have a preference for the stimulus epoch (left) and during the ITI (right). Blue bars indicate current trial variables, yellow bars indicate previous trial variables.

**Supplementary Figure 14**

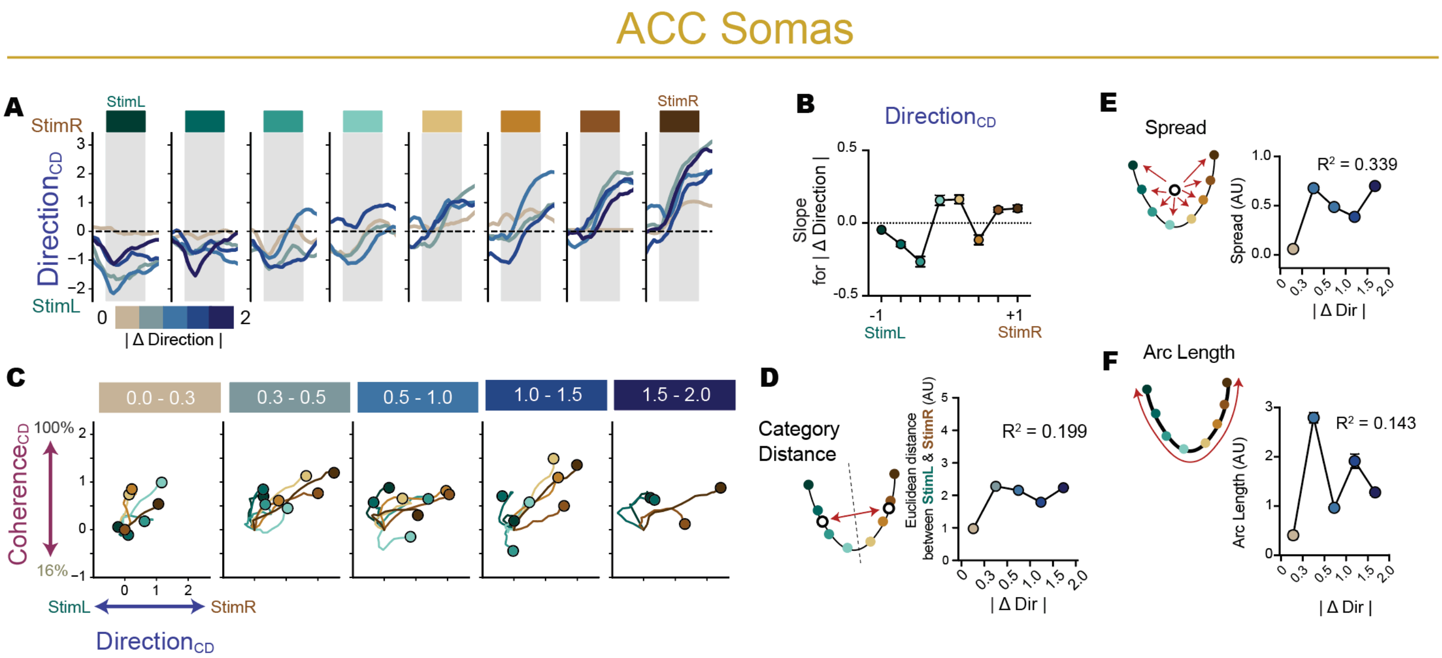


**Supplementary Figure 14: ACC population geometry does not exhibit systematic |ΔDir|-dependent expansion**

(**A**) Trial-averaged ACC population projections onto the DirectionCD for each of the 8 unique stimuli, separated by |ΔDir| (absolute change in stimulus direction from the previous trial). (**B**) Quantification of projection shifts along DirectionCD as a function of |ΔDir|. Modulation as quantified by slopes for |ΔDir| for each stimulus. Slope derived from least squares fit for |ΔDir| to mean projection. Unlike LP-ACC axons, the relationship of stimulus projections to |ΔDir| does not show monotonic scaling. (**C**) Population trajectories projected into the joint DirectionCD-CoherenceCD subspace. Large dots represent each trajectory at the end of the stimulus evaluation period (1.5s after stimulus onset), lines represent their trajectory from baseline (subtracted) upon stimulus onset. (**D** to **F**) Manifold Metrics. (D) Category distance, measured as euclidean distance between mean StimL and StimR representations. (E) Spread, representing dispersion of stimulus representations from centroid, as a function of |ΔDir| (F) Arc length, path distance through ordered signed coherence, as a function of |ΔDir|. Unlike with LP-ACC populations, there was no systematic relationship. $R^2$ reported is a mean derived from linear regression of the metric as a function of |ΔDir| for each cross-validated iteration (n=100 iterations).

**Supplementary Figure 15**

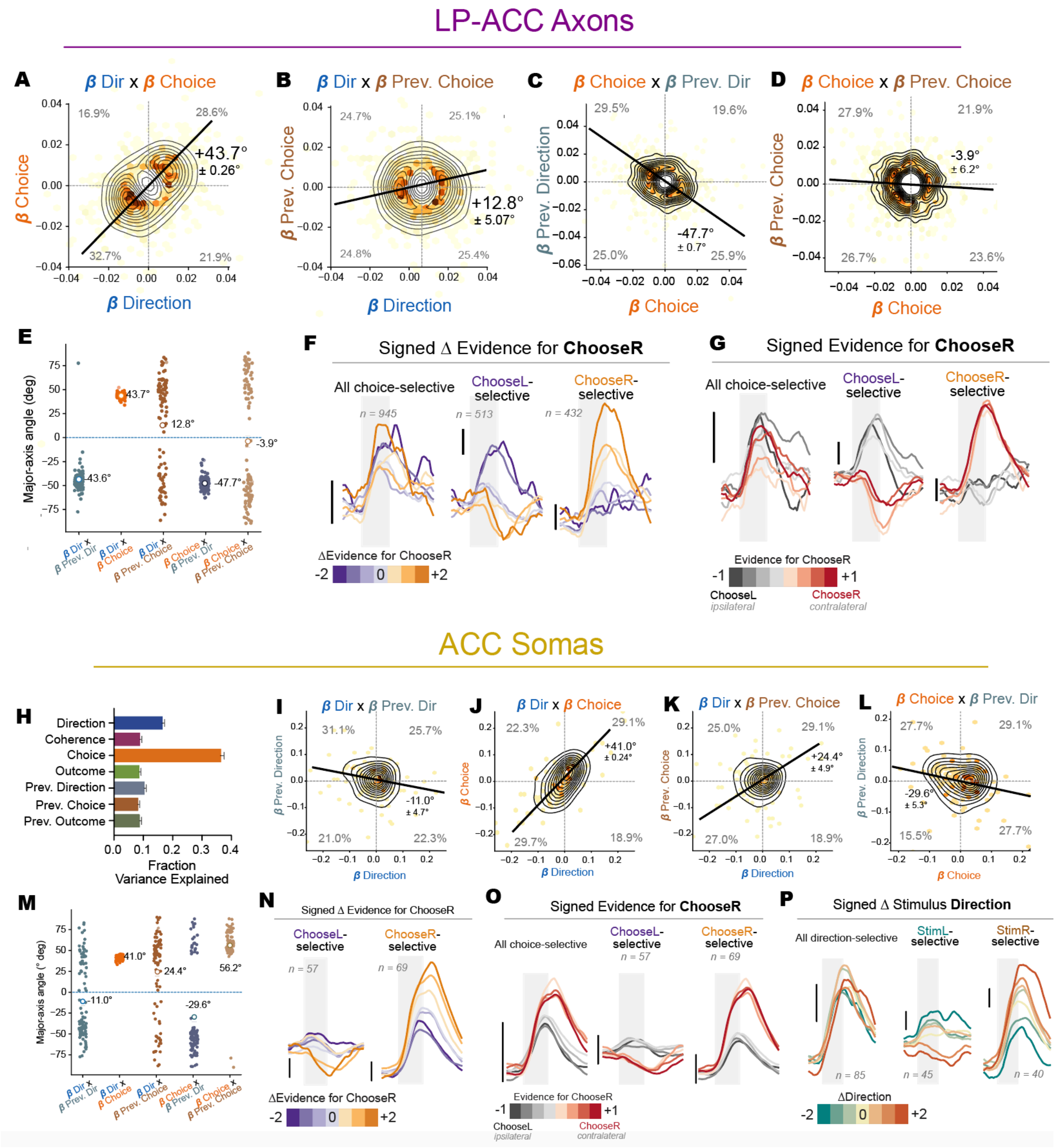

**Supplementary Figure 15: Covariance structure of task-variable weights and subpopulation encoding of current-previous trial interaction in LP-ACC and ACC**

(**A** to **D**) Pairwise correlations between LP-ACC TDR weights for current-trial and history-related variables (A) current stimulus direction (β Dir) vs choice (β Choice), (B) current stimulus direction (β Dir) vs previous choice (β Prev. Choice), (C) current choice (β Choice) vs previous direction (β Prev. Direction), (D) current choice (β Choice) vs previous choice (β Prev. Choice). (**E**) Major axis angle which measures the orientation of the dominant covariance axis between paired regression weights for LP-ACC populations. Each point represents a cross-validated iteration of TDR, with high spread across iterations indicating unstable encoding. (**F**) Subpopulation average of all choice-selective (left), ChooseL-selective (middle) and ChooseR-selective (right) LP-ACC axons, stratified by binned signed Δ Evidence for ChooseR (Current Choice Signed Coherence - Previous Choice Signed Coherence). Choice-selective LP-ACC axons show relatively weak graded modulation. (**G**) Subpopulation average of all choice-selective (left), ChooseL-selective (middle) and ChooseR-selective (right) LP-ACC axons, stratified by binned signed Evidence for ChooseR (only current trial). Choice encoding is more categorical than graded with respect to current evidence. Vertical scale bars for LP-ACC axons all correspond to 0.1 z-scored dFF. (**H**) Fraction of total variance explained by each task CD in ACC neuronal populations derived from targeted dimensionality reduction. ChoiceCD accounts for a larger proportion of population variance than stimulus-direction or history-related CDs. (**I** to **L**) Same pairwise weight correlations as in (A–D), shown for ACC neurons. (I) current stimulus direction (β Dir) vs previous direction (β PrevDir), (J) current stimulus direction (β Dir) vs choice (β Choice), (K) current stimulus direction (β Dir) vs previous choice (β Prev. Choice), (L) current choice (β Choice) vs previous direction (β Prev. Direction). (**M**) Same as (E) but for ACC neurons. (**N**) Same as (F) but for ACC neurons. Accompanies averages for all choice-selective neurons shown in Fig. 5L. ACC neurons exhibit stronger categorical and contralateral (ChooseR) preference compared to LP–ACC axons. (**O**) ACC choice-selective neurons stratified by signed current evidence, showing robust categorical choice encoding. Vertical scale bars correspond to 0.2 z-scored ΔF/F. Vertical scale bars for ACC neurons all correspond to 0.2 z-scored dFF. (**P**) Subpopulation average of all direction selective (left), StimL- (middle) and StimR-selective (right) ACC neurons, stratified by binned signed ΔDir (Current Signed Coherence - Previous Signed Coherence), showing modulation by trial-to-trial differences in signed stimulus evidence.

**Supplementary Figure 16**

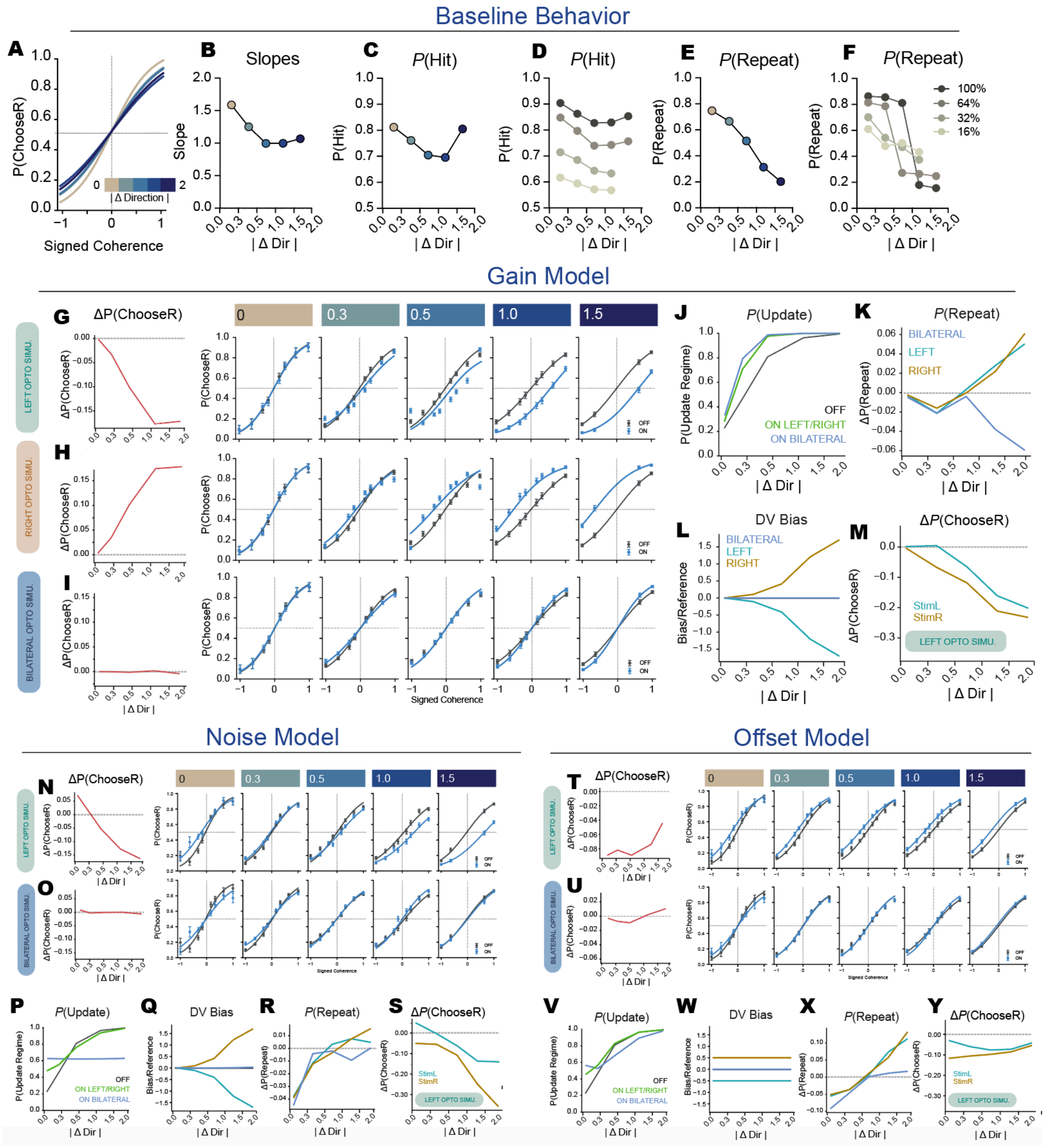

**Supplementary Figure 16: Model validation and comparison of optogenetic implementations**

(**A** to **F**) Validation of the model for reproducing key |ΔDir|-dependent behavioral metrics. Simulations were run through empirical stimulus order statistics (13777 trials, 250 iterations). The model reproduces key |ΔDir|-dependent behavioral features: (A) Psychometric function fits, stratified by |ΔDir| (empirical data: Fig. 1I). (B) Psychometric function slopes show U-shaped modulation by |ΔDir| (empirical data: Fig. 1L). (C) Probability of a hit trial also has non-monotonic relationship with |ΔDir| (Empirical: Fig. 1K). (D) P(Hit) as a function of |ΔDir| separated by stimulus coherence. across stimulus direction and coherence (empirical: fig S2F). (E) P(Repeat) decreases as a function of |ΔDir| (empirical: Fig 1G). (F) P(Repeat) as a function of |ΔDir| plotted separately for different stimulus coherences (empirical: fig S2E). (**G** to **M**) Gain-based optogenetic model. (G to I) Change in P(ChooseR) for a simulated gain-based optogenetic effect (ON - OFF) (left). Psychometric functions (blue, ON and black, OFF) for (G) Left Opto, (H) Right Opto and (I) Bilateral Opto. (J) Probability of being in an update regime (P(Update)) computed for trials based on |ΔDir|. Both unilateral and bilateral optogenetic stimulation increases entry into the Update regime. Note that an update regime is a latent variable that does not determine if an animal switches choice, but determines a new DV computation. (K) Simulations with the gain-based variant, P(Repeat) increases with |ΔDir| in unilateral stimulation, in spite of being in the P(Update) regime, because the choice bias results in reduced choice variability. (L) The bias term which enters the DV to bias choice grows with |ΔDir| as mismatch between |Δd|R and |Δd|L increases due to gain on one |Δd| on only one side. This mismatch is not present in bilateral stimulation, leading to biases canceling out. (M) Unilateral (left) optogenetic stimulation induced change in P(ChooseR) as a function of |ΔDir| stratified by stimulus direction. (**N** to **S**) Noise-based optogenetic model (Δ|Δd| replaced by noise). (N) ΔP(ChooseR) scales with |ΔDir| but exhibits opposite-direction bias at low |ΔDir|. (O) Bilateral stimulation increases Update probability and impairs performance at low |ΔDir|. Remaining panels analogous to (G–M). (**T** to **Y**) Offset-based optogenetic model (constant bias added to |Δd|). (T) Produces ipsilateral bias that does not scale with |ΔDir|, with strongest effects at low |ΔDir|. Remaining panels analogous to (G–M). Simulated right-hemisphere stimulation shows mirror symmetry (not shown).

**Supplementary Figure 17**

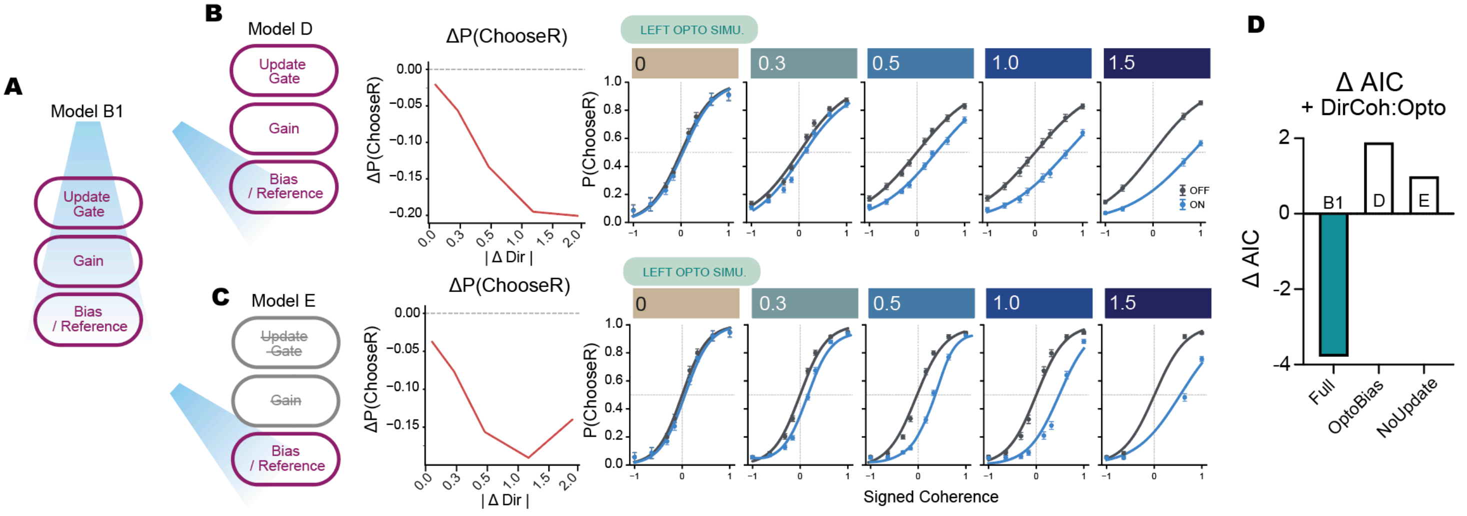


**Supplementary Figure 17: Mechanistic specificity of optogenetic modulation**
(**A**) Schematic of the full mechanistic model (B1), in which optogenetic perturbation modifies LP-derived |ΔDir| signals upstream, thereby influencing all three downstream computations (Update, Gain, and Bias). Behavioral outputs corresponding to this model are shown in Fig. S16G-M. (**B**) Model D ("OptoBias"). Left: Schematic illustrating selective optogenetic modulation of the imbalance (Bias) term while update probability and gain remain determined by true |ΔDir|. Middle: Simulated change in choice probability (ΔP(ChooseR), ON - OFF) under left stimulation, plotted as a function of |ΔDir|. Right: Psychometric functions (ON, blue; OFF, black) shown separately for each |ΔDir| bin. Selective modulation of the bias term reproduces |ΔDir|-dependent lateralized biases while preserving baseline regime structure. (**C**) Model E ("No-Switch"). Left: Schematic in which |ΔDir| enters only through the bias term, with no |ΔDir|-dependent update regime or gain modulation. Middle: Simulated ΔP(ChooseR) under left stimulation. Right: Psychometric functions for each |ΔDir| bin. Although baseline slopes are comparable across |ΔDir| levels, asymmetric perturbation of the bias term alone remains sufficient to generate |ΔDir|-dependent lateralized choice biases. (**D**) GLMM-based model comparison on simulated data. ΔAIC values reflect improvement from adding a DirCoh:Opto term relative to a model containing only |ΔDir|:OptoSide. Only the full mechanistic model (B1) recapitulates simultaneous support for both DirCoh:Opto and |ΔDir|:OptoSide interactions observed in empirical data.

**Table S1:**

| Figure | Dependent variable | Predictor of interest | Estimate (± SE( | Test statistic (p-val) | nMice | nTrials | Dataset |
|---|---|---|---|---|---|---|---|
| 1G | Repeat | AbsDiff | -2.056 ± 0.032 | <2e-16 *** | 22 | 26692 | Baseline (PrevHits) |
| 1K | Hit | AbsDiff | -0.319 ± 0.033 | < 2e-16 *** | 22 | 26692 | Baseline (PrevHits) |
| 2D | Hit | Opto | -0.17304 ± 0.05178 | 0.000833 *** | 6 | 9405 | Left Opto |
| 2H | Hit | Opto | -0.16769 ± 0.09018 | 0.063 | 3 | 3385 | Right Opto |
| 2M | ChooseR | Opto : AbsDiff | -0.279 ± 0.067 | 3.47e-05 *** | 6 | 7102 | Left Opto (PrevHits) |
| 2N | ChooseR | Opto : AbsDiff | 0.242526 ± 0.113795 | 0.03107 * | 3 | 2493 | Right Opto (PrevHits) |
| 2O | ChooseOptoSide | Opto : AbsDiff | 0.265 ± 0.057 | 3.16e-06 *** | 9 | 10115 | Unilateral Opto (PrevHits) |

**Table S1. Statistical details for binomial generalized linear mixed-effects models (GLMMs) reported in main figures.** All dependent variables are binary 0 to 1. AbsDiff (|ΔDir|) are continuous values in the models ranging from 0 to 2. Opto is binary variable. For Hit and Repeat models, stimulus coherence (0.16, 0.32, 0.64, 1.00) was always added as a covariate to control for its influence. All models included (1 | Mouse) as a random effect.

**Table S2:**

| Figure | Dependent variable | Predictor of interest | Estimate (± SE) | Test statistic (p-val) | nMice | nTrials | |
|---|---|---|---|---|---|---|---|
| S1B | ChooseR | Difference | 0.288 ± 0.038 | 7.59e-14 *** | 22 | 35460 | All baseline |
| | | Difference : PreviousOutcome | -0.422 ± 0.038 | < 2e-16 *** | | | |
| S1E | Repeat | AbsDiff | 2.099 ± 0.070 | <2e-16 *** | 22 | 8768 | All baseline (PrevError) |
| S1F | Hit | AbsDiff | 0.901 ± 0.070 | <2e-16 *** | | | |
| S1M | Omit | AbsDiff | 0.121 ± 0.038 | 0.0016 ** | 22 | 40214 | All baseline + Omissions |
| S2B | ChooseR | PrevChoice | 0.703 ± 0.069 | < 2e-16 *** | 22 | 26692 | All baseline (PrevHits) |
| | | AbsDiff | 0.202 ± 0.051 | 8.58e-05 *** | | | |
| | | PrevChoice : AbsDiff | -0.366 ± 0.085 | 1.84e-05 *** | | | |
| S2C | Repeat | PrevChoice | -0.309 ± 0.121 | 0.0106 * | | | |
| | | AbsDiff | -2.110 ± 0.043 | < 2e-16 *** | | | |
| | | PrevChoice : AbsDiff | 0.098 ± 0.060 | 0.1011 | | | |
| S2E | Repeat | DotsCoh | 2.406 ± 0.098 | < 2e-16 *** | | | |
| | | AbsDiff | -0.877 ± 0.069 | < 2e-16 *** | | | |
| | | DotsCoh : AbsDiff | -1.889 ± 0.103 | < 2e-16 *** | | | |
| S2F | Hit | DotsCoh | 1.920 ± 0.102 | < 2e-16 *** | | | |
| | | AbsDiff | -0.286 ± 0.068 | 2.99e-05 | | | |
| | | DotsCoh : AbsDiff | -0.052 ± 0.098 | 0.593 | | | |
| S3B | Omit | Opto | 0.093 ± 0.062 | 0.135 | 9 | 16091 | Unilateral Opto + Omissions |
| S3C | Omit | Opto : AbsDiff | 0.319 ± 0.179 | 0.074127 | 9 | 10182 | Unilateral Opto + Omissions (PrevHits) |

| | | | | | | | |
|---|---|---|---|---|---|---|---|
| S3D | Repeat | Opto | 0.056 ± 0.038 | 0.143 | 9 | 13777 | Unilateral Opto (PrevHits) |
| S3E | Repeat | Opto : AbsDiff | 0.278 ± 0.103 | 0.00692 ** | | | |
| S3F | ChooseR | Opto | -0.203 ± 0.061 | 9.76e-04 *** | 6 | 7102 | Left Opto (PrevHits) |
| S3G | ChooseR | Opto : AbsDiff : StimDir | -0.097 ± 0.240 | 0.685 | | | |
| S3I | ChooseR | Opto | 0.227 ± 0.111 | 0.0417 * | 3 | 2493 | Right Opto (PrevHits) |
| S3J | ChooseR | Opto : AbsDiff : StimDir | -0.263 ± 0.451 | 0.560 | 3 | 3013 | Right Opto (PrevHits) |
| S3N | ChooseR | Opto | -0.264 ± 0.116 | 0.0225 * | 6 | 2303 | Left Opto (PrevErrors) |
| S3P | ChooseR | Opto | -0.177 ± 0.177 | 0.319165 | 3 | 1112 | Right Opto (PrevErrors) |
| S3Q | Hit | Opto | -0.010 ± 0.092 | 0.9057 | 9 | 4321 | Unilateral Opto (PrevErrors) |
| S4B | Hit | Opto | 0.109 ± 0.107 | 0.306 | 3 | 2358 | Bilateral Opto (PrevHits) |
| | Hit | Opto | -0.041 ± 0.089 | 0.644 | 3 | 3285 | Bilateral Opto |
| S4G | Omit | Opto | -0.433 ± 0.218 | 0.0468 * | 3 | 3417 | Bilateral Opto + Omissions |
| | Omit | Opto : AbsDiff | -0.040 ± 0.552 | 0.941 | 3 | 2445 | Bilateral Opto + Omissions (PrevHits) |

| | | | | | | | |
|---|---|---|---|---|---|---|---|
| S4I | ChooseR | Opto | -0.038 ± 0.107 | 0.722 | 3 | 2358 | Bilateral Opto (PrevHits) |
| S4J | ChooseR | Opto : AbsDiff | 0.029 ± 0.116 | 0.797 | | | |
| S4K | ChooseR | Opto : AbsDiff : StimDir | -0.337 ± 0.341 | 0.323 | | | |
| S5C | Hit | Opto | -0.026 ± 0.076 | 0.732 | 3 | 4485 | 593nm Left Opto |
| S5D | ChooseR | Opto | 0.108 ± 0.087 | 0.215 | 3 | 3461 | 593nm Left Opto (PrevHits) |
| S5E | Omit | Opto | 0.027 ± 0.154 | 0.862 | 3 | 4689 | 593nm Left Opto |
| S5F | ChooseR | Opto : AbsDiff | 0.0819 ± 0.198 | 0.679 | 3 | 3461 | 593nm Left Opto (PrevHits) |
| S5G | ChooseR | Opto : AbsDiff : StimDir | -0.319 ± 0.391 | 0.4150 | | | |
| S5H | Hit | Opto : AbsDiff | -0.141 ± 0.183 | 0.442 | | | |
| S5I | Hit | Opto : AbsDiff : StimDir | 0.159 ± 0.372 | 0.669 | | | |
| S6B | Hit | Opto | 0.0641 ± 0.0939 | 0.495 | 3 | 3306 | Left V1 Opto |
| S6G | Omit | Opto | 0.179 ± 0.176 | 0.307 | 3 | 3532 | Left V1 Opto (+ Omissions) |
| S6I | ChooseR | Opto | -0.0114 ± 0.123 | 0.926 | 3 | 2208 | Left V1 Opto (PrevHits) |
| S6J | ChooseR | Opto : AbsDiff | -0.135 ± 0.262 | 0.606 | | | |
| S6K | ChooseR | Opto : AbsDiff : StimDir | -0.498 ± 0.518 | 0.336 | | | |
| S7C | ChooseR | PrevOpto | -0.0226 ± 0.0712 | 0.751 | 6 | 5005 | Left Opto (PrevHits + OptoOFF) |
| S7D | Hits | PrevOpto | -0.118 ± 0.070 | 0.093 | | | |
| S7E | Repeat | PrevOpto | -0.0942 ± 0.0594 | 0.113 | | | |
| S7G | ChooseR | PrevOpto | 0.0779 ± 0.1222 | 0.524 | 6 | 1687 | Left Opto (PrevError + OptoOFF) |
| S7H | Hits | PrevOpto | 0.0233 ± 0.1203 | 0.846 | | | |
| S7I | Repeat | PrevOpto | 0.00556 ± 0.103 | 0.957 | | | |
| S7J | ChooseR | PrevOpto : AbsDiff | -0.0354 ± 0.153 | 0.817 | 6 | 5005 | Left Opto (PrevHits |

| | | | | | | | |
|---|---|---|---|---|---|---|---|
| S7K | ChooseR | PrevOpto : AbsDiff : StimDir | -0.107 ± 0.303 | 0.724 | | | + OptoOFF ) |
| S7L | Hit | PrevOpto : AbsDiff | -0.0385 ± 0.145 | 0.791 | | | |
| S7M | Repeat | PrevOpto : AbsDiff | 0.235 ± 0.149 | 0.114 | | | |
| S8C | ChooseR | PrevOpto | 0.0319 ± 0.0606 | 0.597 | 6 | 7651 | Left Reinforcement Epoch Opto (PrevHits) |
| S8D | Hits | PrevOpto | 0.0803 ± 0.0611 | 0.189 | | | |
| S8E | Repeat | PrevOpto | -0.0427 ± 0.0499 | 0.393 | | | |
| S8G | ChooseR | PrevOpto | 0.0513 ± 0.1283 | 0.689 | 6 | 1876 | Left Reinforcement Epoch Opto (PrevError) |
| S8H | Hits | PrevOpto | 0.0513 ± 0.1271 | 0.686 | | | |
| S8I | Repeat | PrevOpto | -0.101 ± 0.105 | 0.335 | | | |
| S8J | ChooseR | PrevOpto : AbsDiff | 0.0668 ± 0.1412 | 0.636 | 6 | 7651 | Left Reinforcement Epoch Opto (PrevHits) |
| S8K | ChooseR | PrevOpto : AbsDiff : StimDir | -0.0617 ± 0.2821 | 0.827 | | | |
| S8M | Hit | PrevOpto : AbsDiff | -0.0246 ± 0.1344 | 0.854 | | | |
| S8N | Repeat | PrevOpto : AbsDiff | -0.0969 ± 0.1386 | 0.485 | | | |

**Table S2. Statistical details for binomial generalized linear mixed-effects models (GLMMs) reported in supplementary figures.** All dependent variables are binary 0 to 1. Predictor AbsDiff (|ΔDir|) enters the model as continuous values in the models ranging from 0 to 2. Predictor "Difference" (ΔDir) has continuous values ranging from -2 to 2. PreviousOutcome, Opto, StimDir , PreviousChoice and PrevOpto were all binary variables. For Hit, Repeat and Omit models, stimulus coherence (0.16, 0.32, 0.64, 1.00) was always added as a covariate to control for its influence. All ChooseR models included variables from our best fit base model (Fig. 1N, Table S3 Model D) in addition to our predictor of interest. All models included (1 | Mouse) as a random effect.

**Table S3:**
**(A)**

| | | Formula | nPar | AIC | BIC | LL | Comparison | LRT $\chi^2$ | Df | pVal |
|---|---|---|---|---|---|---|---|---|---|---|
| **A** | **Base** | ChooseR ~ DirCoh | 3 | 28800 | 28824 | -14397 | | | | |
| **B** | **PrevChoice** | ChooseR ~ DirCoh + **PrevChoice** | 4 | 28586 | 28618 | 14289 | B vs A | 216 | 1 | < 2.2e-16 *** |
| **C** | **X \|ΔDir\|** | ChooseR ~ DirCoh + **PrevChoice*\|ΔDir\|** | 6 | 28570 | 28620 | -14279 | C vs B | 19.022 | 2 | 7.402e-05 *** |
| **D** | **X \|ΔDir\| + \|ΔDir\|-Gain** | ChooseR ~ DirCoh + **DirCoh*\|ΔDir\|** + **PrevChoice*\|ΔDir\|** | 7 | 28536 | 28593 | -14261 | D vs C | 36.605 | 1 | 1.446e-09 *** |
| **D** | **PrevDir** | ChooseR ~ DirCoh + **PrevDirCoh** | 4 | 28685 | 28717 | -14338 | D vs A | 116.9 | 1 | < 2.2e-16 *** |
| **E** | **PrevChoice*DotsCoh** | ChooseR ~ DirCoh + **PrevChoice*DotsCoh** | 6 | 28572 | 28621 | -14280 | E vs B | 18.001 | 2 | 0.0001233 *** |

**(B)**

| | Estimate | Std. Error | Z | Pr(>\|z\|) | |
|---|---|---|---|---|---|
| **(Intercept)** | -0.5585 | 0.07523 | -7.425 | 1.13e-13 | *** |
| **DirCoh** | 2.28097 | 0.06164 | 37.006 | $< 2 \times 10^{-16}$ | *** |
| **AbsDiff** | 0.35918 | 0.05688 | 6.314 | 2.71e-10 | *** |
| **PreviousChoiceR** | 0.80418 | 0.07120 | 11.294 | < 2e-16 | *** |
| **DirCoh:AbsDiff** | -0.38146 | 0.06258 | -6.095 | 1.09e-09 | *** |
| **AbsDiff:PreviousChoiceR** | -0.66019 | 0.09696 | -6.809 | 9.84e-12 | *** |

**Table S3: Model comparison and selected GLMM coefficients for |ΔDir|-dependent choice behavior (related to Fig. 1N).** (**A**) Comparison of candidate binomial generalized linear mixed-effects models (GLMMs) predicting P(ChooseR), demonstrating improved fit with inclusion of |ΔDir| terms. (**B**) Fixed-effect coefficients for the selected model from (A): *ChooseR ~ DirCoh + DirCoh x |ΔDir| + PrevChoice x |ΔDir| + (1 | Mouse)* (model D). Model fit included N = 26,692 trials from n = 22 mice. Only previously rewarded trials were included in these analyses.

**Table S4:**

**(A)**

| | | nPar | AIC | LL | Comparison | ΔAIC | LRT $\chi^2$ | Df | pVal |
|---|---|---|---|---|---|---|---|---|---|
| **a** | **Base**<br>ChooseR ~ DirCoh + DirCoh:AbsDiff + AbsDiff * PreviousChoice | 7 | 14426 | -7205.9 | | | | | |
| **b** | **+ Side Bias**<br>ChooseR ~ *Base* + **OptoSide** | 8 | 14402 | -7193.0 | b vs a | -24 | 25.772 | 1 | 3.843e-07 *** |
| **c** | **Global Gain**<br>ChooseR ~ *Base* + **DirCoh:Opto** | 8 | 14401 | -7192.4 | c vs a | -25 | 27.052 | 1 | 1.981e-07 *** |
| **d** | **Side-specific Gain + Side bias**<br>ChooseR ~ *Base* + **DirCoh*OptoSide** | 9 | 14398 | -7189.9 | d vs b | -4 | 6.2733 | 1 | 0.01226 * |
| **e** | **PrevChoice-dependent + Side bias**<br>ChooseR ~ *Base* + **PrevChoice*OptoSide** | 9 | 14404 | -7193 | e vs b | +2 | 1e-04 | 1 | 0.9936 |
| **f** | **\|ΔDir\| Bias**<br>ChooseR ~ *Base* + **AbsDiff:OptoSide** | 8 | 14397 | -7190.3 | f vs b | -5 | | | |
| **f'** | **\|ΔDir\| + Side Bias**<br>ChooseR ~ *Base* + **AbsDiff*OptoSide** | 9 | 14399 | -7190.3 | f' vs f | +2 | 0.5813 | 1 | 0.4458 |
| | | | | | f' vs b | -3 | 5.4084 | 1 | 0.02004** |
| **g** | **Global Gain + Side bias**<br>ChooseR ~ *Base* + **DirCoh:Opto** + **OptoSide** | 9 | 14383 | -7182.6 | g vs c | -18 | 19.513 | 1 | 9.992e-06 *** |

| | | | | | | | | | |
|---|---|---|---|---|---|---|---|---|---|
| **h** | **Global Gain + \|ΔDir\| Bias**<br>ChooseR ~ *Base* + **DirCoh:Opto** + **AbsDiff:OptoSide** | 9 | 14380 | -7181.8 | h vs f | -17 | 19.084 | 1 | 1.251e-05 *** |
| | | | | | h vs g | -3 | | - | |
| | | | | | h vs c | -21 | 22.631 | 1 | 1.963e-06 *** |
| **i** | **Global Gain + Side Bias + \|ΔDir\| Bias**<br>ChooseR ~ *Base* + **DirCoh:Opto** + **AbsDiff*OptoSide** | 10 | 14382 | -7180.8 | i vs h | +2 | 0.6221 | 1 | 0.4303 |
| **j** | **Global \|ΔDir\|-dependent gain modulation**<br>ChooseR ~ *Base* + **DirCoh:Opto** + **AbsDiff:OptoSide** + **DirCoh:AbsDiff:Opto** | 10 | 14381 | -7180.6 | j vs h | +1 | 0.8376 | 1 | 0.3601 |
| | ChooseR ~ *Base* + **DirCoh:Opto** + **AbsDiff:OptoSide** + **DirCoh:AbsDiff:OptoSide** | | | | | | | | |
| **k** | **Side-specific, \|ΔDir\|-dependent gain modulation**<br>ChooseR ~ *Base* + **DirCoh:Opto** + **AbsDiff:OptoSide** + **DirCoh:AbsDiff:OptoSide** | 10 | 14382 | -7181.0 | j vs i | +1 | 0.1293 | 1 | 0.7192 |

**B)**

| | **Estimate** | **Std. Error** | **Z** | **Pr(>\|z\|)** | |
|---|---|---|---|---|---|
| **(Intercept)** | -0.5404 | 0.11175 | -4.836 | 1.32e-06 | *** |
| **DirCoh** | 2.56896 | 0.09107 | 28.208 | $< 2 \times 10^{-16}$ | *** |
| **PreviousChoiceR** | 0.87486 | 0.10192 | 8.584 | $< 2 \times 10^{-16}$ | *** |
| **AbsDiff** | 0.33501 | 0.08224 | 4.0746 | 4.62e-05 | *** |
| **DirCoh:AbsDiff** | -0.46394 | 0.08810 | -5.266 | 1.40e-07 | *** |
| **PreviousChoiceR : AbsDiff** | -0.51049 | 0.13944 | -3.661 | 0.000251 | *** |
| **DirCoh : OptoTRUE** | -0.41650 | 0.09331 | -4.464 | 8.06e-06 | *** |
| **AbsDiff : OptoSide** | 0.25249 | 0.05367 | 4.704 | 2.55e-06 | *** |

**Table S4. GLMM model comparison and coefficients for optogenetic perturbation data (related to Fig. 2P).** (**A**) Comparison of candidate binomial generalized linear mixed-effects models (GLMMs) predicting P(ChooseR) under optogenetic stimulation, including terms for stimulation, |ΔDir|, and their interaction. (**B**) Fixed-effect coefficients for the selected model from (A) (Model h): *ChooseR ~ DirCoh + DirCoh:AbsDiff + AbsDiff * PreviousChoice + DirCoh:Opto + AbsDiff:OptoSide*
Model fit included N = 13777 trials from n = 9 mice. All previous hit trials from unilateral LP-ACC stimulation sessions were included.

**Table S5:**

| | Comparison Signal | Opto Mode | Unilateral bias | Unilateral impaired performance with \|ΔDir\| | Bias increases with \|ΔDir\| | Mirror-symmetry for L & R opto | Bilateral cancellation of lateralization | Bilateral preserved performance |
|---|---|---|---|---|---|---|---|---|
| **A1** | **Signed Difference** | Gain | ✗ | - | - | - | - | ~ Improved high \|ΔDir\| |
| **A2** | | Offset | ✓ | ✗ Yes but most impaired low \|ΔDir\|, less impairment with greater \|ΔDir\| | ✗ | ✓ | ✓ | ~ Impaired low \|ΔDir\|, Small improvement intermediate \|ΔDir\| |
| **A3** | | Noise | ✗ | - | - | - | - | ✗ Impairment across \|ΔDir\| |
| **B1** | **Unsigned Comparator** | Gain | ✓ | ✓ | ✓ | ✓ | ✓ | ~ Small improvement at higher \|ΔDir\| |
| **B2** | | Offset | ✓ | ✗ Yes but most impaired low \|ΔDir\|, less impairment with greater \|ΔDir\| | ✗ | ✓ | ✓ | ~ Impaired low \|ΔDir\|, Improved intermediate \|ΔDir\| |
| **B3** | | Noise | ✓ | ✗ Most impaired at low \|ΔDir\| and high \|ΔDir\| | ~ Yes but with contralateral bias at low \|ΔDir\| | ✓ | ✓ | ~ Impaired low \|ΔDir\| |
| **C1** | **Prev Choice Remap** | Gain | ✓ | ✓ | ✓ | ✓ | ✓ | ~ Improvement at higher \|ΔDir\| |
| **C2** | | Offset | ✗ | ✓ | - | - | - | ~ Impaired low \|ΔDir\|, Small improvement intermediate \|ΔDir\| |
| **C3** | | Noise | ✓ | ~ Reduced slope at low \|ΔDir\| without bias | ✓ | ✓ | ✓ | ✗ Impairment across \|ΔDir\| |
| **D** | **Unsigned Comparator (OptoBias)** | Gain | ✓ | ✓ | ✓ | ✓ | ✓ | No change |
| **E** | **Unsigned Comparator (no update)** | Gain | ✓ | ✓ | ✓ | ✓ | ✓ | No change |

**Table S5. Model-based evaluation of candidate mechanisms for optogenetic effects.** Candidate model architectures and perturbation mechanisms were evaluated against empirical constraints derived from unilateral and bilateral LP–ACC stimulation experiments. Rows indicate model variants; columns indicate whether each variant reproduces specific behavioral signatures. Symbols denote qualitative agreement (✓), disagreement (✗), or partial agreement (~). Further elaborated in Supplementary Note 1.
